# Tipping Points and Emergent Phases of Species Diversity in Mutualistic Ecological Networks under Global Warming

**DOI:** 10.64898/2026.02.23.706386

**Authors:** Yongzheng Sun, Jiayi Wang, Man Qi, Guanghui Wen, Wei Lin, Roberto Salguero-Gómez, Philip K. Maini

## Abstract

Global warming poses a severe threat to ecosystem resilience and biodiversity, potentially driving species loss, declining diversity, and ecosystem collapse. Accurately predicting tipping points of ecological resilience under global warming remains a fundamental challenge. Here, we develop a theoretical framework to identify climate-induced tipping points in mutualistic ecological networks, capturing critical transitions under warming scenarios. We show that rising temperatures, increasing intraspecific competition, and deteriorating mutualistic interaction can drive ecosystems through three distinct dynamical phases of species diversity: stable full coexistence, stable partial coexistence, and total extinction. Notably, ecosystems with low species abundances are particularly vulnerable, facing heightened risks of diversity loss and extinction. Moreover, the warming rate modulates these transitions, as slower warming grants endangered species a window of time within which to adjust and attenuate the loss of resilience. Applying this framework to assess various emission mitigation pathways, we find that “pollute first, clean up later” scenarios pose a threat to biodiversity, leading to irreversible biodiversity losses even after subsequent emission reductions. Our findings offer critical insights into the timing and mechanisms of ecological tipping points under climate change and underscore the urgency of proactive mitigation strategies.

---

Climate change and biodiversity loss are among the greatest threats to ecosystem integrity and supply of nature’s contribution to the human species [1]. Global warming is an incontrovertible reality and is acknowledged as a principal driver of profound disruptions to ecosystem structure and function [2–4]. Over the past century, global temperatures have exhibited a marked warming trend, accelerating in recent decades [5]. Notably, the latest report has indicated that, as global warming approaches the tipping point of 1.5°C, the Earth’s climate and natural systems are approaching multiple irreversible tipping points at an unprecedented rate [6]. The combined impacts of ongoing climate change and persistent anthropogenic pressures are driving an increase in extreme climate events [7, 8] and posing acute threats to global diversity [1, 9, 10]. Indeed, evidence indicates irreversible coral reef loss from cumulative heat stress [8] and a two-thirds decline in terrestrial insects by 2100 even for 1.5°C warming [10]. These alarming trends underscore the urgent need to predict species’ responses to ongoing global warming to inform effective ecosystem management and conservation [11].

Faced with global warming and its accompanying extreme weather events, ecosystem structure and function have been severely impaired. Ecosystems differ in their capacity to recover, known as resilience–the ability to restore core structure and function after disturbance–first conceptualized by Holling [12] and now a cornerstone of ecological theory [13–18]. As environmental stress intensifies, ecosystem resilience gradually declines and may eventually reach a tipping point, beyond which abrupt and often irreversible shifts–or even collapse–of ecosystem structure and function can occur [19–21]. Beyond the magnitude of stress, the rate of environmental change can also trigger tipping points. Even when the final stress level remains within the nominal tolerance range, rapid environmental change may outpace the system’s capacity to adapt, leading to a loss of resilience–a phenomenon known as rate-induced tipping [22–24]. Consequently, developing reliable approaches to detect and predict temperature tipping points under accelerating global warming is critical for safeguarding ecosystem stability in rapidly changing environments.

Mutualistic ecological networks are particularly vulnerable to rising temperatures, with key biological processes–such as plant flowering phenology and pollinator foraging rhythms–being markedly affected [25–27]. Temperature modulates mutualistic interactions and competition (both inter- and intra-specific) within these networks [28, 29]. Combined with the complex bidirectional feedbacks between temperature and biotic interactions [30], this can make ecosystems highly susceptible to perturbations that disrupt species balance and elevate extinction risks.

Without considering temperature effects, studies have linked mutualistic network resilience to structural and interaction complexity [31] and to mortality rate [24]. However, predicting temperature-induced tipping points for resilience loss in mutualistic ecological networks is highly challenging due to their high dimensionality, nonlinear dynamics, and structural complexity. Therefore, elucidating how global warming dynamically reshapes the structure and function of mutualistic networks through climate–biotic interactions is crucial for understanding ecosystem resilience [32, 33].

Biodiversity underpins ecosystem function and resilience, and the gradual loss of species can serve as an early-warning signal for biodiversity conservation [34, 35]. However, existing studies mainly focus on predicting tipping points of ecosystem collapse, often overlooking these gradual species losses that precede systemic failure. Considering the long-standing debate over whether diversity enhances ecosystem resilience [36], and given the intrinsic structural complexity of ecosystems and the multidimensional pressures imposed by global warming [1, 9, 10], understanding how these factors shape biodiversity and stability remains a central challenge. Although emergent patterns of species diversity driven by species interactions are well documented [34, 35], it remains unclear whether analogous temperature-induced phase transitions occur. Moreover, in response to rising temperatures, species may maintain resilience through adjustments in intraspecific competition and the strength of mutualistic interactions [30], yet how these mechanisms affect system-level diversity remains unknown.

To address these gaps, we develop a theoretical framework combining network theory with mean-field approximation in temperature-sensitive mutualistic ecological networks. This approach yields analytical predictions for the tipping points governing ecosystem transitions under warming, providing a deeper mechanistic understanding of the resilience of mutualistic networks. We uncover three successive dynamical phases–stable full coexistence, partial coexistence, and total extinction–emerging under rising temperatures, stronger intraspecific competition, and weakened mutualistic interactions. Ecosystems with initially low species abundances are particularly vulnerable, showing rapid resilience loss, while the rate of warming modulates outcomes by allowing slower increases to sustain ecosystem function. Applying the framework to climate mitigation scenarios shows that delayed emission reductions impose severe, often irreversible, diversity risks. Our findings provide new insights into the timing and mechanisms of climate-driven ecosystem collapse and inform strategies for biodiversity conservation under accelerating global change.

## Results

### Prediction of tipping points

To investigate the impact of climate warming on ecosystems, we consider long-term climate warming trends and introduce temperature-dependent saturated functional responses. Due to the prolonged pollination handling time caused by rising temperatures during the annual peak pollination period [29], we incorporate temperature into classical mutualistic ecological networks as in [21]

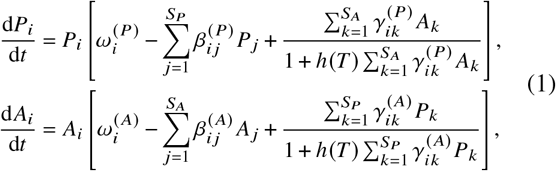

where *P*_*i*_ and *A*_*i*_ denote the abundances of the *i*-th plant and pollinator, respectively; 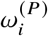 and 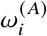 represent their intrinsic growth rates; 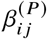 and 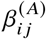 (for *i* ≠ *j*) describe interspecific competition among plants and pollinators; and 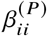 and 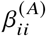 account for intraspecific competition among plants and pollinators, respectively, as illustrated in Fig. 1*A*. Typically, intraspecific competition is stronger than interspecific competition [25], i.e., *β*_*ii*_ ≫ *β*_*i j*_ . Mutualistic interactions may be present or absent, and 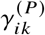 represents the mutualistic interaction from each pollinator species to each plant species, and 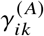 measures the mutualistic interaction between each plant species and each pollinator species. In general, *γ*_*ik*_ depends on the number of mutualistic species *K*_*i*_, namely,

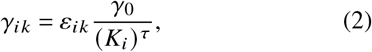

where *γ*_0_ is a constant representing mutualistic strength, if *ε*_*ik*_ = 1, plants and pollinators interact (otherwise, *ε*_*ik*_ = 0), *K*_*i*_ is the number of mutualistic interaction species, and *τ* determines the strength of the tradeoff between the interaction strength and the number of interactions. If there is no tradeoff (*τ* = 0), the network topology will have no effect on the strength of the mutualistic interactions. In contrast, a full tradeoff (*τ* = 1) means that the network topology will affect the strength of the mutualistic interactions.

**Fig. 1.**
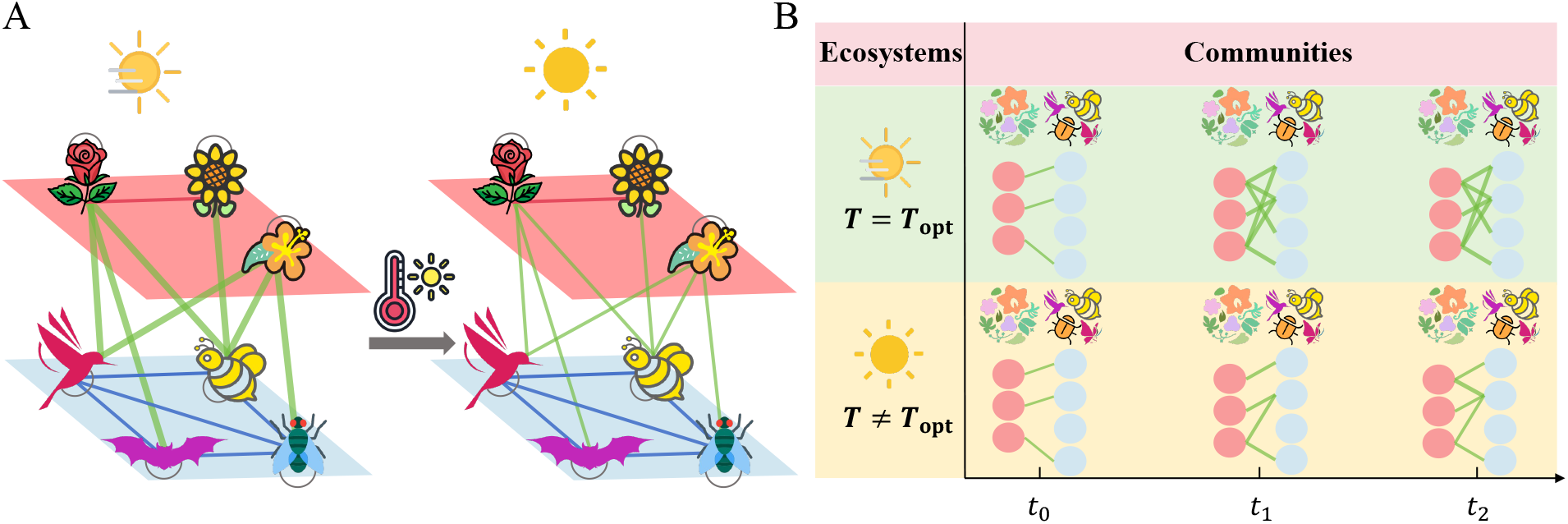
Global warming induces weakening of mutualistic ecological networks described by Eq. 1. (*A*) Schematic illustration of mutualistic networks reveals the negative impact of global warming. Species are represented as nodes on a two-layer network (plants: red layer; pollinators: blue layer), and interactions are shown as links: plant interspecific competition (red), pollinator interspecific competition (blue), intraspecific competition (gray), and mutualistic interactions (green). (*B*) Impact of temperature variation on the formation of mutualistic networks. When temperature deviates from the optimum, increased handling time weakens mutualistic interactions, thereby slowing the formation of mutualistic networks and reducing their structural complexity. Thinner links imply that mutualistic interactions weaken.

The last term on the right-hand side of Eq. **1** represents a saturating mutualistic function: at low partner density, the mutualistic benefit increases approximately linearly, whereas at high partner density, the species becomes “service-saturated”. The function *h* (*T*) quantifies the saturation rate (or handling time), representing a species’ capacity to assimilate the benefits from its partners. Larger *h* accelerates saturation and reduces marginal benefits from additional partners. Temperature affects the handling time required for biological processes such as digestion and pollination, as shown in Fig. 1*B* and Fig. S1 in *SI Appendix*. To capture potential temperature effects on this process, we model *h* as a temperature-dependent parameter representing the handling time of mutualistic interactions [28, 29], which preserves the unimodal relationship as [37]

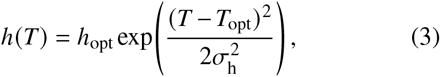

where *h*_opt_ denotes the value of *h* (*T*) at the optimal temperature *T*_opt_, and *σ*_h_ describes the thermal breadth. This formulation implies that *h* (*T*) reaches its minimum at *T*_opt_ and increases as temperature deviates from this optimum, leading to low-density saturation of mutualistic benefits (see Fig. S1*A, SI Appendix*). This assumption enables the analysis of how rising temperatures affect mutualistic ecological networks.

Analyzing the dynamical properties of the mutualistic network in Eq. **1** is challenging due to its high dimensionality and nonlinearity. To facilitate mathematical analysis and numerical computation of mutualistic network resilience under temperature fluctuations, we apply mean-field theory [20, 38] to derive a one-dimensional (1D) model (see Methods section) that integrates the abundances of both plants and pollinators as follows [31]

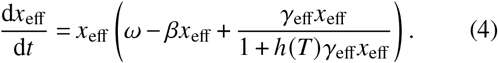

Here, *x*_eff_ denotes the effective average abundance of plants and pollinators; *β* = *β*_eff_ + *β*_0_ characterizes the effective average competition which combines the effects of intraspecific competition *β*_0_ and effective average interspecific competition *β*_eff_. The parameters *ω* and *γ*_eff_ correspond to the effective average intrinsic growth rate and effective average mutualism, respectively. Therefore, we predict that the network of interspecific competition and mutualistic interactions in the microscopic description can be collapsed into the two macroscopic resilience parameters *β*_eff_ and *γ*_eff_. This simplification enables the application of the bifurcation analysis framework–originally developed for 1D systems–to high-dimensional mutualistic networks under temperature influence. This approach enables accurate prediction of system responses to diverse perturbations and precisely identifies the tipping points when resilience is lost.

Ecosystems may experience declines in diversity and even species extinctions as global warming intensifies [1– 4, 7–10]. We next investigate, through theoretical analysis, the tipping points that trigger abrupt species extinctions. In the 1D model (Eq. **4**), with fixed effective average competition and mutualism, the system dynamics are primarily governed by temperature *T*, allowing us to examine how temperature variations influence the resilience of mutualistic networks. As global warming continues to drive temperatures beyond the optimum, we focus our analysis on the regime where *T* > *T*_opt_, guided by the temperature dependence described in Eq. **3**.

We begin by assuming a slowly varying temperature, representing a gradual warming trend. Clearly, the 1D model described by Eq. **4** has a trivial steady state at *x*_eff_ = 0, representing the extinction of all species. The nontrivial equilibria of Eq. **4** are determined by

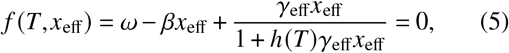

where the function *f* governs the system dynamics. Within a nonlinear-dynamics framework, the climate-warming-induced phase transition—where species diversity collapses from coexistence to extinction—emerges as a bifurcation in Eq. **4**, with temperature *T* acting as the control parameter. Beyond the critical temperature *T*_c_, the system shifts from coexistence to total extinction. Below, we derive the tipping points *T*_c_ that induce the loss of system resilience.

Initial species abundances *x*_0_ critically influence ecological resilience [39]. At sufficiently high initial abundances and temperatures *T* above a critical threshold, *f* (*T, x*_eff_) can always remains negative, rendering *x*_eff_ = 0 the sole stable fixed point of Eq. **4** and driving species extinction. The tipping temperature for extinction is derived below (see Methods)

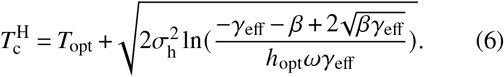

For relatively low initial abundances, an abundance-dependent estimate of the tipping temperature can be obtained (see Methods)

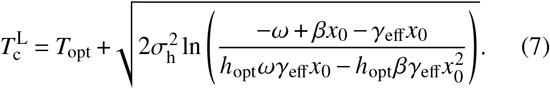

Importantly, 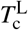 is determined not only by the network structure, effective average competition and mutualism, but also by the initial species abundances. The range of low initial abundances is specified in the Methods.

Figure 2*A* illustrates the stable and unstable states in the 1D model and the tipping points associated with resilience loss. When 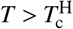, as marked with a pentagram, species in ecosystems with high initial abundances ultimately face extinction. Furthermore, for systems with low initial abundances, extinction occurs at the point 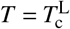, marked with a diamond. Building on the theoretical results, we verify the predicted tipping points of resilience for the empirical ecological network *M*_*PL*_005, sourced from the Web of Life database (www.web-of-life.es). Unless stated otherwise, simulations use high initial abundances *x*_0_ = 2 and low initial abundances *x*_0_ = 0.05. Temperature is varied over *T* ∈ [25, 40] with other parameters held constant. The system described by Eq. **1** is simulated, and the effective average abundance is plotted in Fig. 2*B* for high (green) and low (blue) initial abundances. Comparison between theoretical results (lines) and numerical simulations (symbols) shows that the 1D model (Eq. **4**) accurately predicts tipping points of resilience for the high-dimensional network (Eq. **1**).

**Fig. 2.**
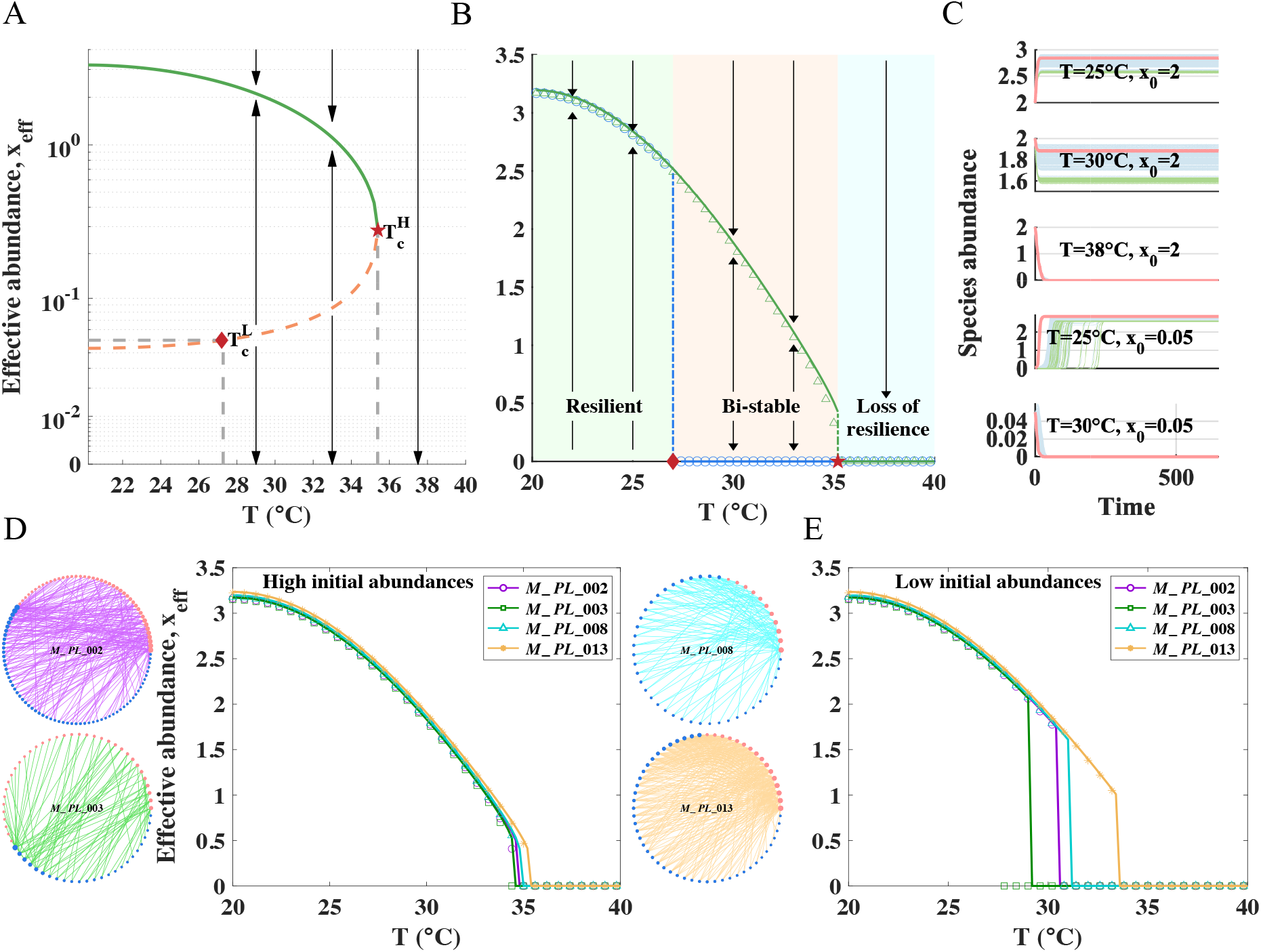
Rising temperatures drive mutualistic networks from coexistence to collapse. (*A*) Phase diagram of the reduced 1D model for the *M*_*PL*_005 ecological network shows a stable high-abundance resilient state, an unstable low-abundance state, and a stable collapsed state. Initial abundances govern ecosystem resilience under warming: ecosystems with high initial abundances remain resilient until 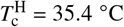, whereas those with low initial abundances reach the tipping point earlier at 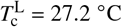. (*B*) The reduced 1D model accurately reproduces the resilience of the original high-dimensional ecosystem (Eq. **1**) under high (green triangles) and low (blue circles) initial abundances, where arrows point to the stable fixed points of the system. Rising temperatures induce transitions from a resilient state to a bi-stable state, and ultimately to a collapsed state. (*C*) Community responses to rising temperatures under high (top three panels) and low (bottom two panels) initial abundances. At *T* = 25°C slightly above the optimal temperature, all species maintain full coexistence. When the temperature rises to 30°C, ecosystems with high initial abundances remain in a stable coexistence state, whereas those with low initial abundances have already collapsed. The blue, green, and red lines show the temporal evolution of pollinators (governed by Eq. **1**), plants (governed by Eq. **1**), and their effective average abundances (governed by Eq. **4**). (*D*) and (*E*) The reduced 1D model (Eq. **4**), broadly applicable across empirical ecological networks, accurately predicts resilience loss in mutualistic networks under (*D*) high and (*E*) low initial abundances. High and low initial abundances correspond to *x*_0_ = 2 and *x*_0_ = 0.05, respectively. Other parameters used are 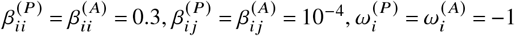, *h*_opt_ = 0.5, *T*_opt_ = 20, *σ*_h_ = 15, *τ* = 0.5, and *γ*_0_ = 12.

Figure 2*B* shows that when the temperature surpasses the optimum by approximately 15°C (high initial abundances) or 7°C (low initial abundances), even a slight further increase triggers a dramatic change in system dynamics. The temperature *T* governs system resilience, partitioning the functional space into three distinct regions. For low initial abundances, the resilient state (green, 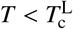) has a single stable positive equilibrium *x*_eff_ > 0. In the bi-stable state (red, 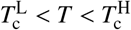), recovery depends on the initial condition. In the non-resilient state (blue, 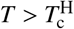), only the extinction state *x*_eff_ = 0 is stable, and recovery is impossible. To illustrate the effects of temperature and initial abundances on species states, we present abundance evolution diagrams in Fig. 2*C*. These show the temporal changes of pollinators (blue), plants (green), and the effective average abundance (red) under conditions corresponding to Fig. 2*B*. Systems with lower initial abundances are particularly prone to extinction and highly sensitive to temperature changes.

To further demonstrate the effectiveness of the theoretical prediction of tipping points and understand the influence of initial abundances on the system, we conduct a deeper investigation into the impact of initial abundances on tipping points across empirical networks, as shown in Fig. 2 *D* and *E*. By comparing Fig. 2 *D* and *E*, we observe that under global warming, while the overall loss of resilience remains limited, species with low initial abundances experience a more rapid decline in resilience, in line with previous theoretical finding [39]. Mathematically, this pattern arises from the coexistence of a stable attractor and an unstable equilibrium. Communities with initially low abundances near the unstable equilibrium are more prone to resilience loss [31, 40]. Ecologically, the existence of an alternative steady state as temperature rises implies a small stochastic perturbation in species abundances can result in a shift in stable state. To identify which network structures are more vulnerable to global warming, we simulate ecosystems with well-defined topologies—Scale-Free networks, Erdős–Rényi networks, and Watts–Strogatz small-world networks [38] including interspecific competition network and random bipartite graphs for mutualistic networks (see Fig. S2 in the SI Appendix). We find that tipping points are only weakly affected by interspecific competition network topology, but are strongly mediated by mutualistic connectivity, especially in systems with low initial species abundances.

### Emergent phases of species diversity

Global warming not only causes species extinction but also drives the loss of species diversity [1, 9, 10]. To examine how temperature influences species diversity, we analyze the fraction of persisting species at stable equilibrium as temperature increases. For high initial abundances, simulations of the system described by Eq. **1** show that rising temperatures drive the system from stable full coexistence (SFC, phase I) to stable partial coexistence (SPC, phase II), and ultimately to total extinction (TE, phase III), as shown in Fig. 3*A*. Figure 3*B* plots the survival fraction against temperature, clearly displaying the sequence of dynamical phases. Unlike ecosystems with high initial abundances, the results in Fig. 3 *C* and *D* suggest that systems with low initial abundances exhibit only two distinct phases (I and III), highlighting their heightened vulnerability and propensity for abrupt collapse. In particular, the phase I–III and II–III transitions are accurately predicted by the analytical expressions for the tipping temperature identified in Fig. 3 (pentagrams and diamonds). These results align with recent empirical studies [1, 9, 10] demonstrating that climate warming drives species diversity loss, supporting the robustness of our findings.

**Fig. 3.**
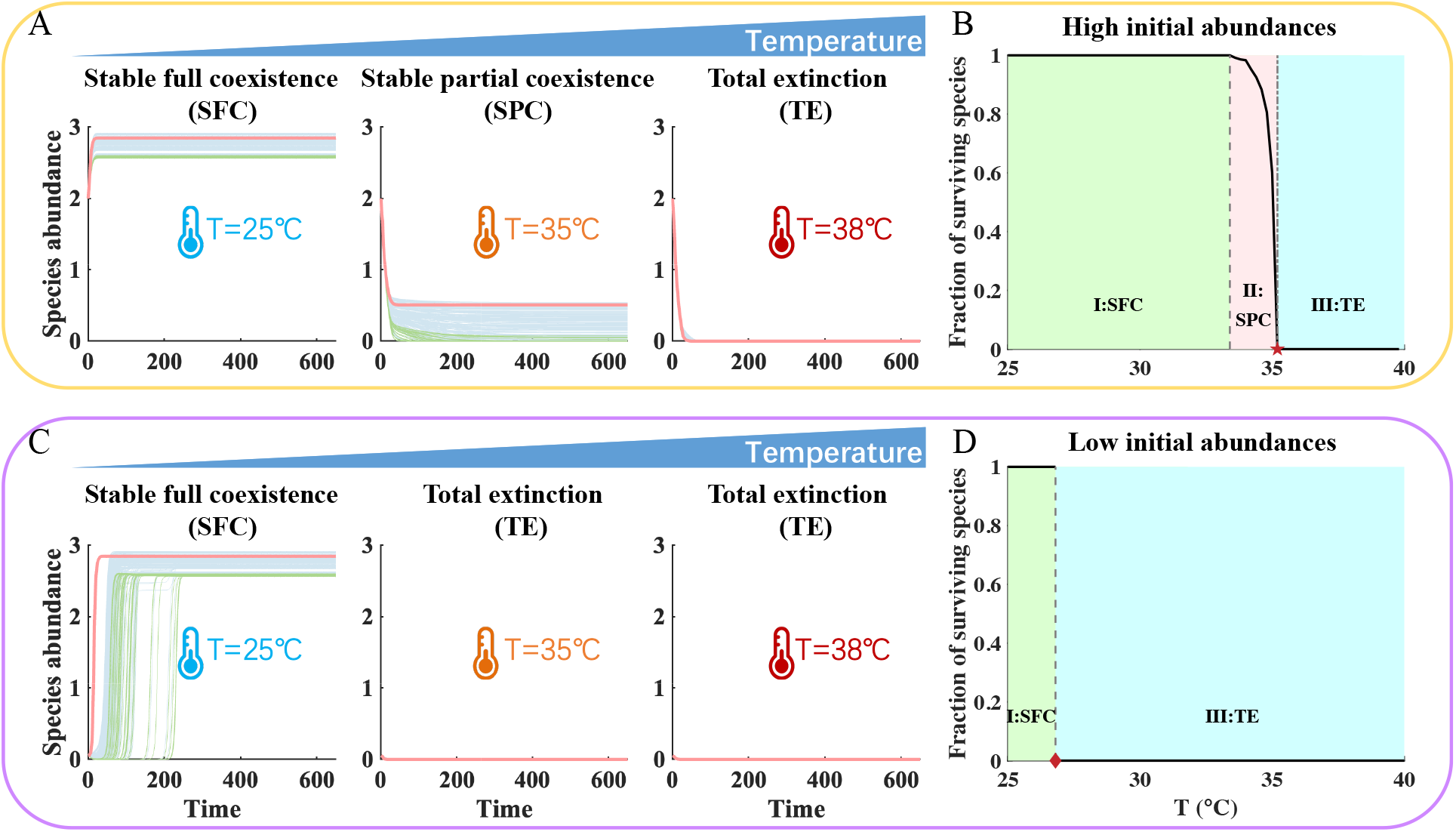
Emergent phase transitions in species diversity under rising temperatures. (*A*) Change of species abundances for pollinators (blue), plants (green), and their effective average abundance (red) under high initial abundances across different environmental temperatures. Communities transition from stable full coexistence at *T* = 25°C, to partial coexistence at *T* = 35°C, and ultimately to total extinction at *T* = 38°C. (*B*) Mapping the survival fraction against temperature reveals that rising temperatures can induce a sequence of phase transitions from phase I to phase II and then to phase III. (*C*) Unlike ecosystems with high initial abundances, low-initial-abundance systems do not exhibit a transient partial coexistence as temperature rises from 25°C to 38°C, showing only stable full coexistence at *T* = 25°C and total extinction at *T* = 35°C and *T* = 38°C. (*D*) Low-initial-abundance systems are more vulnerable to global warming, with the phase transition from Phase I to Phase III occurring earlier as temperature increases. The network is *M*_*PL*_005, with all other parameters the same as in Fig. 2.

### Influence of species interactions and buffering effect

As global warming progresses, not only do species abundances and species diversity respond, but interaction types and strengths also shift as a result of ecological adaptation, species dispersal and migration [41, 42]. Here, we investigate how variations in mutualism strength, intraspecific competition, and interspecific competition links affect system resilience along temperature gradients, as shown in Fig. S3 in the *SI Appendix*. Our results show that increasing mutualism strength, reducing intraspecific competition and interspecific competition links enhance species abundances and resilience to non-optimal temperatures. However, deviations from optimal temperature constrain the range of competition and mutualism strengths that keep the system stable [30]. These findings demonstrate that species interactions and temperature jointly regulate the whole ecosystem resilience.

Given that interspecific competition exerts a relatively weak influence on mutualistic networks [25], we primarily focus on investigating how two key factors–intraspecific competition and mutualistic interaction strength–affect the resilience and emergent phases of species diversity. We gradually increase the intraspecific competition *β*_0_ ∈ [0.3, 2.1] and the mutualistic strength *γ*_0_ ∈ [2, 11], as illustrated in Fig. 4 *A-D*. Ecosystems characterized by strong mutualistic strength and weak intraspecific competition exhibit higher tipping points, indicating greater tolerance to global warming. Figure 4 *B* and *C* illustrate emergent phases defined by the species survival fraction, under varying temperatures and interactions (intraspecific competition and mutualistic strength), respectively. The emergent phases of diversity are jointly regulated by temperature and species interactions, with the critical thresholds of different emergent phases indicated by gray dashed lines. Specifically, at a fixed temperature, increasing intraspecific competition while maintaining a constant mutualistic interaction strength drives the ecosystem from phase I to phase III, reflecting substantial species loss. In contrast, with intraspecific competition held constant, enhancing the mutualistic interaction strength shifts the system from phase III back to phase I, demonstrating that stronger mutualistic strength can effectively restore species diversity. The corresponding tipping point 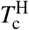 of the system described by Eq. **1** under high initial abundances is presented in Fig. 4*D*. These results support a recent theory according to which species survive by increasing mutualistic interaction strength and decreasing intraspecific competition [31].

**Fig. 4.**
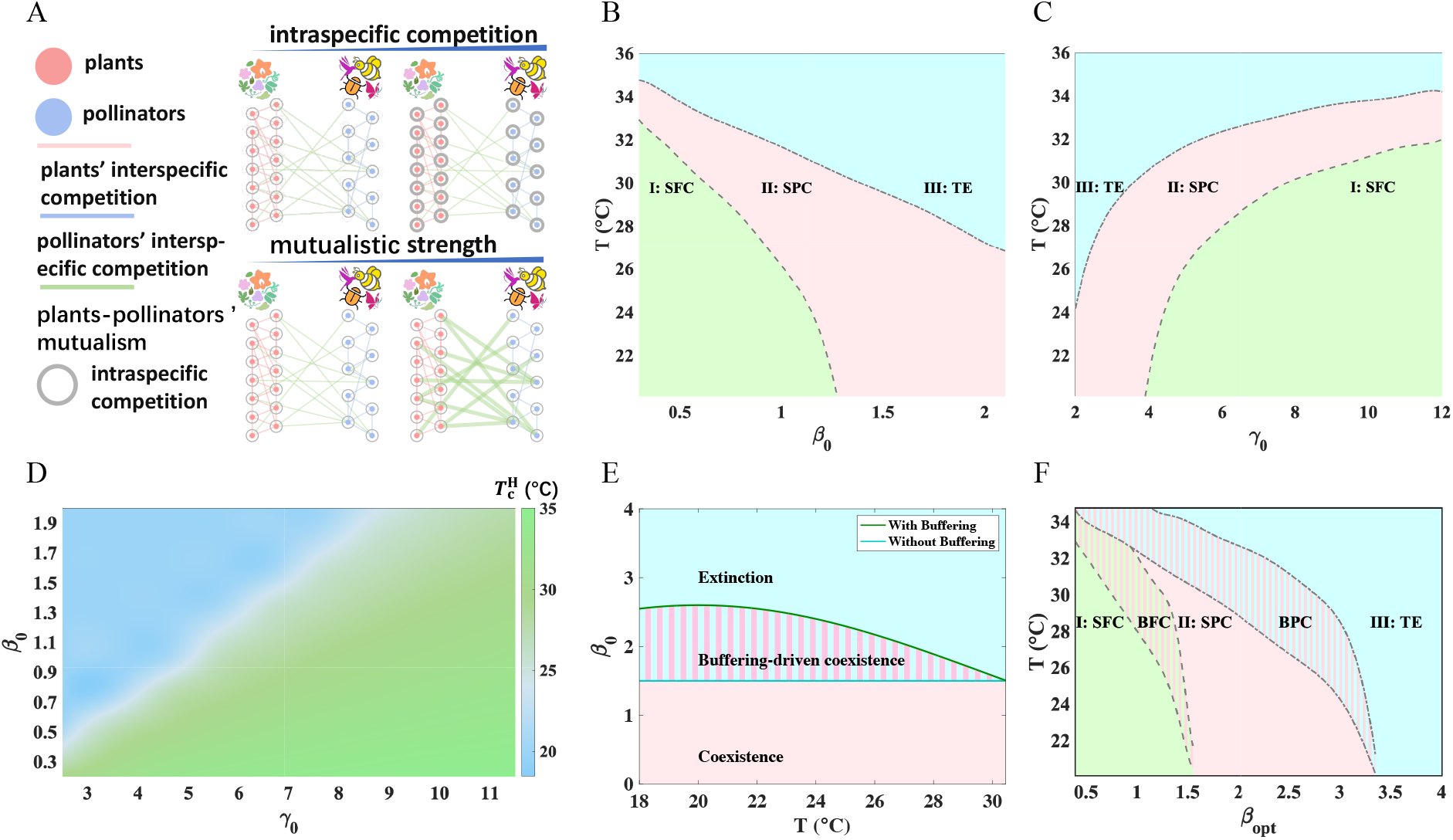
Stronger mutualistic interaction and weaker intraspecific competition significantly delay the onset of resilience loss, and the buffering of intraspecific competition enhances ecosystem resilience to warming. (*A*) Complex mutualistic interaction and intraspecific competition within and between plants and pollinators shape the resilience of mutualistic ecosystems under global warming. (B) Increasing the strength of intraspecific competition while keeping the mutualistic interaction fixed (*γ*_0_ = 10) leads to a continuous decline in diversity, with the system transitioning from full coexistence (phase I) to partial coexistence (phase II), and ultimately to extinction (phase III). (*C*) Enhanced mutualistic interactions promote the recovery of species diversity. For a fixed intraspecific competition (*β*_0_ = 0.6), increasing the strength of mutualistic interactions can restore species diversity, resulting in a transition from phase III back to phase I. (*D*) Stronger mutualistic interactions and weaker intraspecific competition increase the tipping temperature and delay resilience loss under global warming. (*E*) Compared with the level of species diversity in the absence of buffering of intraspecific competition, buffering responses that modulate intraspecific competition via the unimodal thermal dependence in Eq. **8** expand the temperature range for coexistence and delay community collapse. (*F*) Buffering of intraspecific competition generates a buffering-driven full coexistence phase (BFC) between SFC and SPC, and a buffering-driven partial coexistence phase (BPC) between SPC and TE, which delays species loss and stabilizes community diversity under environmental stress. In (*B*)-(*F*), high initial abundances are used. In (*E*)-(*F*), the adjustment rate parameter is set to *s* = 10. All other parameters are the same as in Fig. 2.

Facing the environmental stress imposed by global warming, species often deploy adaptive strategies to preserve ecosystem diversity. The Demographic Buffering Hypothesis suggests that populations may buffer key demographic rates against environmental fluctuations, thereby reducing the risk of collapse [43]. The Stress-Gradient Hypothesis posits that as environmental stress intensifies, competition may weaken and even shift toward mutualism [41]. Together, these hypotheses indicate that species can actively adjust their traits and interactions to mitigate the impacts of environmental stress and preserve ecosystem stability. Here, since interspecific competition exerts a relatively weak influence on mutualistic networks [25], we consider the buffering mechanism of intraspecific competition. Building on these perspectives, we further assume that intraspecific competition exhibits a unimodal thermal dependence within a mechanistic framework incorporating life-history trade-offs [44, 45], reflecting the highest resource requirement and competition intensity at optimal temperature

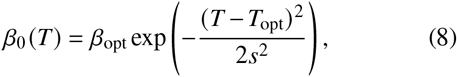

where *β*_opt_ denotes the maximum intensity of intraspecific competition at the optimal reproductive temperature *T*_opt_, and *s* reflects the rate at which competition declines as temperature deviates from the optimum (as shown in Fig. 4*E*). Figure 4*F* illustrates the emergent species diversity phases driven by rising temperatures and the unimodal thermal dependence of intraspecific competition. Comparing the simulations in Fig. 4*F* with those in Fig. 4*B*, we find that increasing temperatures and stronger intraspecific competition can still induce transitions from SFC to TE. However, when species can actively respond to thermal stress by reducing intraspecific competition, the buffering-driven full coexistence (BFC) phase and partial coexistence (BPC) phase emerge, which significantly mitigates the risk of species loss. Consequently, as temperature deviates from the optimal value, declining intraspecific competition delays the tipping points of species extinction (see Fig. S4 in *SI Appendix*). Our findings also demonstrate that tropical species are highly vulnerable to extinction due to climate change (see Fig. S5 in *SI Appendix*).

### Rate-induced tipping and emergent phases

Accelerated global warming, fueled by human activities, has become a critical ecological concern [7, 8]. The warming rate over the past two centuries exceeds any period in history [5], raising urgent questions about impacts on ecosystem stability and biodiversity. Here, we focus on how rapid warming rates, rather than absolute temperatures alone, can trigger tipping points and emergent phases in species diversity.

How rate-induced tipping emerges in the system is revealed through basin-of-attraction theoretical analysis of the 1D model given by Eq. **4** (see Methods). The function *f* (*T, x*_eff_) can not only reflect the stability of the system but also determine the range of the basin of attraction. For a fixed temperature *T* ∈ [25, 32.2]°C, Fig. 5*A* illustrates the dependence of *f* (*T, x*_eff_) on the effective average abundance *x*_eff_. Each curve of *f* (*T, x*_eff_) intersects the x-axis at two points. The intersection point on the right (solid circle) corresponds to the stable equilibrium state, while the intersection on the left (open circle) corresponds to the unstable equilibrium state. Specifically, at 25°C, when the effective average abundance lies to the left of the green dashed line, the ecosystem exhibits a negative gradient, resulting in a gradual decline in species abundances that ultimately leads to extinction. Conversely, when the effective average abundance is to the right of the green dashed line, the system can self-regulate and converge to a stable equilibrium state (solid circle). Therefore, for any given temperature *T*, there exist two basins of attraction corresponding to survival and extinction. The boundary between these basins at *T* = 32.2°C is indicated by the red dashed line. It is evident that as the temperature deviates further from the optimal temperature, the extinction basin of attraction progressively expands. This pattern indicates that, when the warming rate accelerates and the initial abundances of endangered species cannot adjust rapidly enough, the system trajectory may enter the extinction basin of attraction, ultimately resulting in species extinction. Our analysis serves as a dynamical foundation for intuitively understanding the subsequent simulations.

**Fig. 5.**
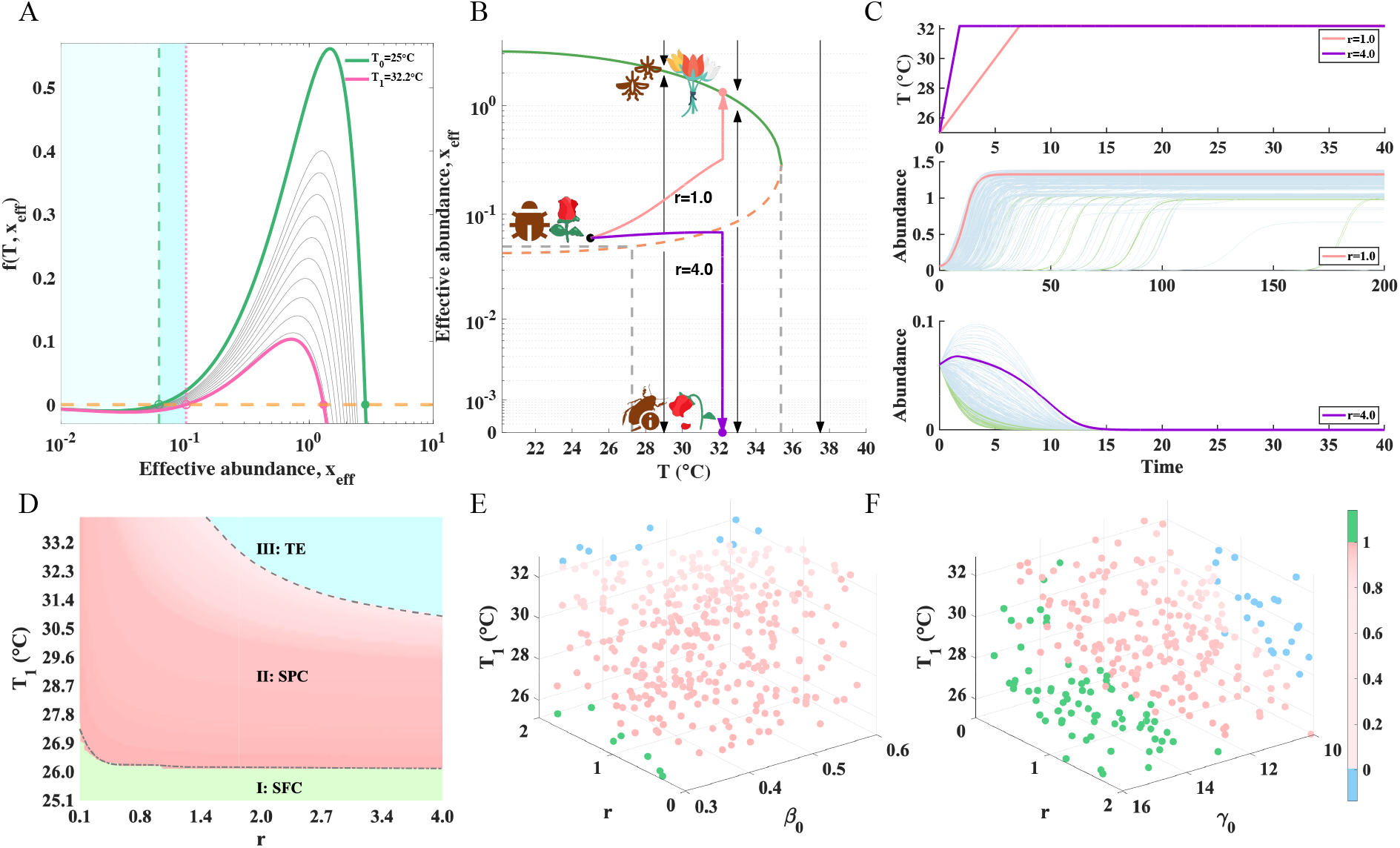
Rate-induced tipping and emergent phase transitions of diversity demonstrate that rapid warming can induce ecological collapse. (*A*) Boundaries of extinction basin of attraction for *T* ∈ [25, 32.2]°C (green dashed line: *T*_0_ = 25°C; red dashed line: *T*_1_ = 32.2 C) show that the extinction basin of attraction (light to dark blue) expands with rising temperature. The green solid line (*T*_0_ = 25°C), red solid line (*T*_1_ = 32.2°C), and black solid line represent the system gradient described by Eq. **4**, with their intersections with the orange dashed line indicating zero-rate equilibria. (*B*) The resilience function of the reduced 1D model (Eq. **4**) shows that ecosystems starting near low initial abundances rapidly approach collapse under fast warming (*r* = 4, purple), whereas controlling the warming rate (*r* = 1, red) allows the system to stabilize at a high-abundance steady state via adjustment. (*C*) Temperature variation under *r* = 1 and *r* = 4 (upper panel), as well as the time series of pollinators (blue), plants (green) in system described by Eq. **1** and effective average abundance (red and purple) in 1D model (Eq. **4**) at *r* = 1 (middle panel) and *r* = 4 (lower panel). (*D*) For fixed intraspecific competition (*γ*_0_ = 12) and mutualistic strength (*β*_0_ = 0.4), as the global warming rate increases, communities lose species (transition from phase I to phase II) before eventual collapse (phase III). (*E*) and (*F*) Fractions of species surviving under varying temperatures and warming rates: (*E*) fixed mutualistic strength (*γ*_0_ = 12) with varying intraspecific competition, and (*F*) fixed intraspecific competition (*β*_0_ = 0.4) with varying mutualistic strength. Increasing mutualistic strength or reducing intraspecific competition promotes species coexistence. The network is *M*_*PL*_005, with all other parameters the same as in Fig. 2.

To investigate how warming rate affects ecosystem dynamics, we consider a scenario in which temperature increases linearly over time from *T*_0_ = 25°C to *T*_1_ = 32.2°C at a constant rate *r*, described by

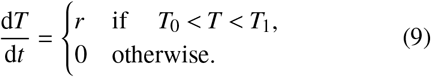

To better demonstrate rate-induced species extinction, we perform simulations initiated from low initial abundances (*x*_0_ = 0.06), recognizing that resilience loss is highly sensitive to initial conditions [46]. Results for slow (*r* = 1) and fast (*r* = 4) warming rates are shown in Fig. 5 *B* and *C*. It can be observed that within the same temperature variation range (*T*_0_ to *T*_1_), species subjected to a slow warming rate ultimately maintain high abundance levels, whereas under a fast warming rate, they decline into extinction. A slow warming rate provides endangered species with a critical time window to adjust and recover, allowing their abundances to increase to high levels and restoring their resilience despite rising temperatures. Our theoretical results indicate that an accelerated rate of warming undermines ecosystem resilience by accumulating “ecological debt” [8, 47]. This finding is consistent with empirical evidence from coral reef systems, where sustained thermal stress constrains biological regeneration–leading to the accumulation of such ecological debt–and drives the progressive degradation and increased collapse risk of reef habitats under global warming [8]. Therefore, measures aimed at mitigating the rate of global warming are of paramount importance for maintaining ecosystem resilience.

Next, we explore the impacts of warming rates (*r*) and the final temperature state (*T*_1_) on the emergent phases of species diversity. The species survival fraction is plotted over the *r*–*T*_1_ parameter space, as shown in Fig. 5*D*. It can be observed that the system transitions from phase I to phase II, and subsequently to phase III, as either *r* or *T*_1_ increases. In Fig. 5 *E* and *F*, we comprehensively examine the combined effects of mutualistic strength (*γ*_0_), intraspecific competition without an buffering mechanism (*β*_0_), warming rate (*r*), and final temperature (*T*_1_) on the species survival fraction. The analysis reveals that higher mutualistic strength, weaker competition, lower warming rate, and lower final temperature collectively promote greater diversity. These findings are consistent with the conclusions drawn from Fig. 4 *B* and *C*.

### Ecological role of mitigation pathways

In response to the accelerating impacts of global warming on ecosystems, governments worldwide have implemented various mitigation initiatives, most notably the Paris Agreement [48], which aims to limit the rise in global temperature to well below 2°C above pre-industrial levels, and preferably to 1.5°C. To support these goals, the scientific community has developed the Shared Socioeconomic Pathways (SSPs) framework, which posits that future surface temperature increases are influenced not only by natural systems but also by human socioeconomic activities and development choices. The SSPs comprise five baseline scenarios–SSP1 (Sustainability–Taking the Green Road), SSP2 (Middle of the Road), SSP3 (Regional Rivalry–A Rocky Road), SSP4 (Inequality–A Road Divided), and SSP5 (Fossil-fueled Development–Taking the Highway) [49]. Each is characterized by fundamentally different levels of adaptation capacity and mitigation feasibility, with detailed information provided in Table 1 in *SI Appendix*. In these scenarios, higher radiative forcing corresponds to greater global temperature rise. Therefore, understanding how SSPs influence species status is critical for assessing the impact of future warming on ecological tipping points and emergent phases.

We now investigate the influence of different SSPs on the ecological system resilience. Building upon our previous finding that species in tropical regions are at risk of extinction, we select three representative low-latitude mutualistic networks, as illustrated in Fig. 6*A*. Based on their geographic locations, we acquire temperature projections from multiple climate models under various SSP scenarios via the Earth System Grid Federation (https://esgf-node.llnl.gov/search/cmip6/), and we compute annual average temperatures across months of peak pollination activity. Using these data, we simulate Eq. **1** under conditions of high initial abundances and derive time series of effective average abundance to 2100 for each network and SSP, as shown in Fig. 6 *B–D*. Corresponding temperature projections are also presented. The shaded regions for effective average abundance and temperature represent uncertainties arising from the ensemble of climate model projections. Our theoretical tipping temperature predicted by Eq. **6** (marked by red dashed lines in insets) aligns closely with the simulated onset of species extinction. Notably, under scenarios characterized by unrestrained anthropogenic impacts–such as SSP3-7.0 and SSP5-8.5–ecological systems exhibit a pronounced risk of resilience loss. These findings highlight the critical need for proactive and effective measures to safeguard the stability of the ecosystem.

**Fig. 6.**
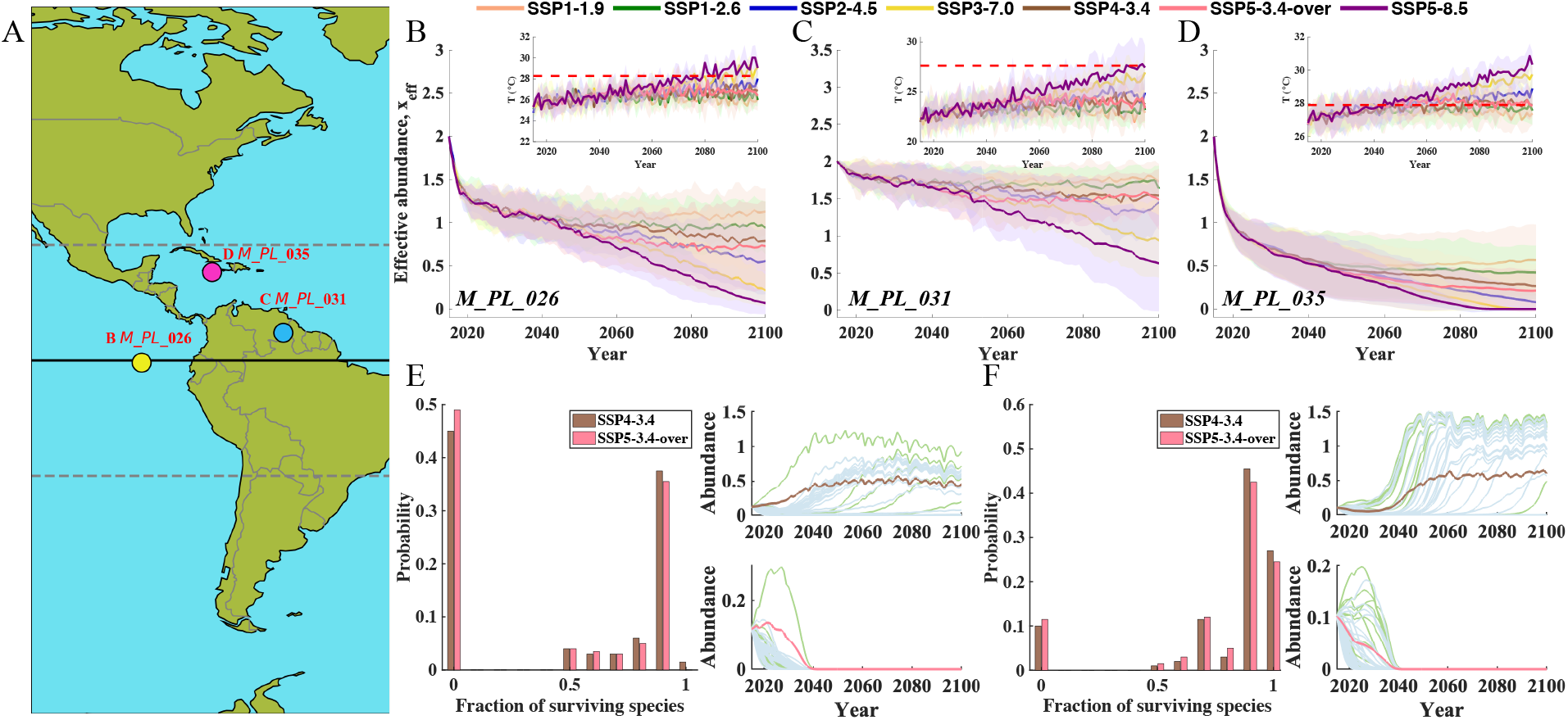
Ecosystem collapse risk intensifies under high-emission Shared Socioeconomic Pathways (SSPs). (*A*) Geographic distribution of empirical mutualistic networks. (*B*)-(*D*) As the temperature increases, ecological systems face a higher risk of resilience loss. Numerical simulation of effective average species abundances under different SSPs are shown, with shaded regions indicating uncertainties from the ensemble of climate model projections, for (*B*) *M*_*PL*_026, (*C*) *M*_*PL*_031, and (*D*) *M*_*PL*_035. Insets: temperature rise trends (solid lines with shaded bands) and tipping point (red dashed lines) predicted by Eq. **6** under different SSPs. (*E*) and (*F*) Faster warming increases the risk of species loss in (*E*) *M*_*PL*_026 and (*F*) *M*_*PL*_031. Left panels show projected survival probabilities under different scenarios by 2100 (*x*_0_ ∈ [0.1, 0.15] for *M*_*PL*_026; *x*_0_ ∈ [0.09, 0.14] for *M*_*PL*_031), while right panels illustrate abundance dynamics for representative initial conditions (*x*_0_ = 0.118 for *M*_*PL*_026; *x*_0_ = 0.103 for *M*_*PL*_031). Here, *h*_opt_ = 0.6, *σ*_h_ = 10, the other parameters are same as in Fig. 2.

Recent studies have highlighted that the over-reliance of climate scientists and the media on worst-case scenarios–often treated as “business-as-usual”–is misleading, and that more realistic scenarios should be used as reference baselines for climate risk assessment [50]. However, even if future greenhouse gas emissions are effectively controlled, does this necessarily imply that ecological resilience will revert to previous levels? To investigate this question, we simulate the fraction of surviving species under two SSP scenarios featuring different warming rates. The results in Fig. 6 *B* and *C* confirm that, under the SSP4-3.4 and SSP5-3.4-over scenarios, temperatures in both the *M*_*PL*_026 and *M*_*PL*_031 networks consistently remain below their tipping points of resilience loss by 2100, enabling analysis of warming rate effects on resilience. In contrast, network *M*_*PL*_035 collapses under these scenarios, making rate effects indistinguishable. Accordingly, we perform simulations using these two SSP scenarios in the respective networks. These two scenarios share similar levels of radiative forcing, implying comparable warming magnitudes. However, SSP4-3.4 relies on phased gradual emission reductions, whereas SSP5-3.4-over represents a “pollute first, clean up later” development model. This model is heavily dependent on large-scale geoengineering to achieve drastic emission cuts in its later stages. Since the occurrence of resilience loss is strongly dependent on the system’s initial conditions [46], we conduct simulations starting from different low initial abundances and calculate the probability of species survival fractions in 2100, as shown in Fig. 6 *E* and *F*. Under the SSP5-3.4-over scenario, the fraction of surviving species is consistently lower than under SSP4-3.4, indicating a greater threat to species diversity.

Crucially, our findings demonstrate that the “pollute first, clean up later” paradigm is fundamentally untenable. As research shows, when the response rate of ecosystems lags behind the pace of climate forcing, consecutive extreme heat events produce cumulative effects, leading to a “mismatch” between ecosystems and climatic conditions. Without sufficient short-term recovery, this mismatch accumulates as “ecological debt”, ultimately lowering the threshold for collapse and making ecosystems more prone to irreversible decline [47]. Furthermore, time-lagged demographic effects–including delayed population recovery and extinction cascades–can decouple environmental improvement from diversity rebound, resulting in a protracted loss of ecosystem function. Together, these insights underscore that the timing of mitigation is as critical as its magnitude: delayed action risks locking ecosystems into degraded states with reduced capacity to recover, even if climatic conditions later stabilize [51].

## Discussion

Tipping phenomena and emergent phases of species diversity are hallmark features of complex ecosystems [13– 15, 34, 35]. Global warming is proceeding at an unprecedented rate, posing severe threats to ecosystem resilience, biodiversity, and the services they provide [1, 7–10]. Predicting the tipping points induced by global warming, at which ecosystems lose resilience and biodiversity, is essential for guiding effective mitigation and conservation strategies. Here, we focus on forecasting the tipping points for resilience loss in mutualistic networks. As temperature rises, the handling time increases, mutualistic interactions between plants and pollinators weaken, driving ecosystems toward collapse, while intraspecific and interspecific competition jointly modulate this process. By mapping high-dimensional, nonlinear ecological dynamics onto a low-dimensional space, our framework captures the essential features of empirical mutualistic networks and accurately predicts the tipping points of resilience loss. We find that as temperature increases, competition intensifies or mutualistic interactions weaken, mutualistic ecosystems transition sequentially from full species coexistence to partial coexistence, and ultimately to collapse, consistent with theoretical predictions of emergent community behaviors. Moreover, the rate of warming regulates these transitions: slower warming provides endangered species with a crucial window to adjust, thereby attenuating the reduction of resilience and maintaining biodiversity. Applying this framework to assess current emission mitigation pathways, we find that “pollute first, clean up later” scenarios pose a fundamental threat to biodiversity, leading to irreversible losses even after subsequent emission reductions.

To assess the potential impact of global warming on the resilience and diversity of mutualistic ecological networks, we model the handling time *h* as a unimodal function of temperature, centered at the optimal temperature *T*_opt_. Although this simplifying assumption neglects possible asymmetries in *h*, i.e., the differential weakening of mutualistic interactions under cooling versus warming [52], it captures the experimentally supported pattern that warming weakens mutualistic interactions: as temperature exceeds the optimum, interaction frequency declines and visit duration shortens [37]. This framework thus provides a parsimonious means to explore how global warming may compromise the stability and functioning of mutualistic networks and impair ecosystem service delivery.

Our findings indicate that initial species abundances play a pivotal role in ecosystem resilience under global warming. Species with low initial abundances face higher extinction risk, triggering cascading diversity and functional losses. This highlights the importance of monitoring baseline population sizes alongside species richness to identify vulnerable species in time. Conservation strategies maintaining or enhancing species abundances can help ecosystems against climate-driven collapse.

Faced with the challenges of global warming and increasingly frequent extreme climatic events, adaptive interactions within and across species can mediate system stability to environmental variability. At the demographic level, populations may adaptively buffer environmental variability by reducing fluctuations in key life-history parameters, thereby stabilizing population growth–a core assumption of the Demographic Buffering Hypothesis [43]. Under warming, such buffering may manifest as decreased intraspecific competition, mitigating the escalating effects of temperature fluctuations from individuals to populations. However, studies have shown that population buffering is not unlimited and may diminish as physical stress intensifies [53]. Once buffering capacity declines, population growth rate begins to decrease. Therefore, incorporating a stress-dependent intrinsic population growth rate will be important in future studies to better explore scenarios under which population buffering collapses.

Our model does not explicitly consider life-history trade-offs among competing species, lateral migration that may create new interactions, or spatial mismatches between plants and pollinators. This simplification allows us to focus on the general effects of interspecific interactions on network stability, providing a basis for incorporating more ecosystem-specific dynamics in future case studies. Our results show that the stability of a mutualistic network is less sensitive to variation in interspecific interaction strength than to changes in intraspecific interaction or mutualism strength, although decreasing interspecific competition can promote species coexistence and enhance network stability, offering a testable hypothesis for future empirical research.

Although enhancing mutualistic interaction strength and reducing intraspecific competition can raise the tipping temperature, as shown in Fig. 4, this capacity is finite: once warming surpasses the new tipping point, mutualistic interaction strength declines and intraspecific competition intensifies, accelerating diversity loss. The limited adaptation of intraspecific competition and mutualistic interactions highlights the prediction that the interplay between external stress intensity and intrinsic reconfiguration may confer resilience under moderate warming, but still leads to collapse once its threshold is exceeded. Future work integrating population models with long-term plant–pollinator time series could empirically test whether such mechanisms fail near climate-driven ecological tipping points.

Our theoretical analysis indicates that both the magnitude and the warming rate of temperature can impact mutualistic networks, potentially triggering system collapses. To reduce the risk of resilience loss, it is crucial to continuously monitor species interactions and slow the pace of climate change, allowing species time to adapt. Our findings not only reveal the key mechanisms by which temperature changes affect ecological network stability but also provide theoretical insights and practical guidance for preserving ecosystem resilience and informing adaptive management strategies under accelerating global warming.

## Methods

### Mean-field theory

To gain deeper insights into tipping points, we conduct a mean-field theoretical analysis. For the original highdimensional network described in Eq. **1**, we assume that the intrinsic growth rates and intraspecific competition coefficients are identical across nodes [21, 31]. Namely, 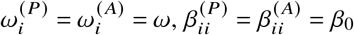. We obtain

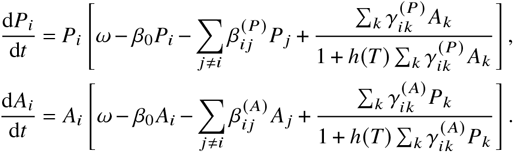

For networks with weak degree correlations, the system’s effective state is quantified by the average nearest-neighbour activity, reflecting the total influence of all neighbours through the node’s degree and the corresponding average nearest-neighbour value [38]. By defining 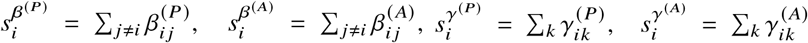, and introducing ℒ(*M*, **y)** = (**1**^⊤^*M***y)/ (1**^⊤^*M***1)** as the influence contributed by the average nearest neighbor nodes, we quantify the total interaction strengths for each node. Then, we apply this operator to both sides of Eq. **1**, respectively. Further we let 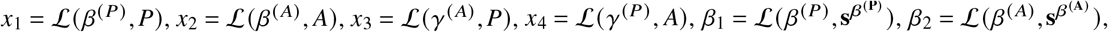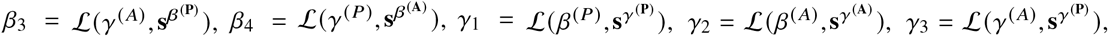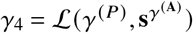. Then

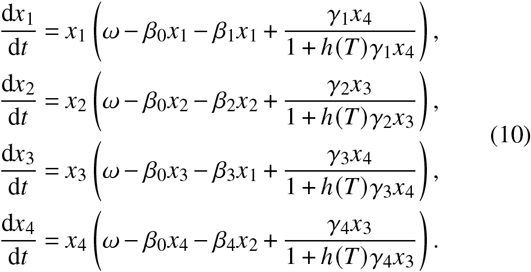

Here, *x*_1_ and *x*_2_ represent the weighted effects of interspecific competition on the abundances of plants and pollinators, respectively; *x*_3_ and *x*_4_ denote the weighted average effects of pollinators and plants on plant abundance *P* and pollinator abundance *A* through mutualistic interactions, respectively. The above analyses allow us to compress the high-dimensional space into a four-dimensional one. To further map the system onto a one-dimensional space, the four-dimensional information is first reconstructed into an adjacency matrix as follows

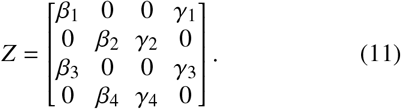

Similarly, we can take *Z* as the input matrix *M* of the operator ℒ(*M*, **y)** in Eq. **10**. Finally, we obtain the effective 1D model, as shown in Eq. **4** of the main text

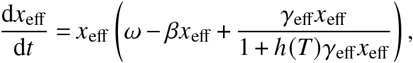

where *β* = *β*_eff_ + *β*_0_, and

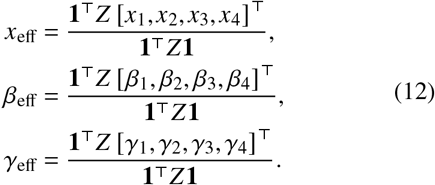

### Calculation of tipping points

At the steady state d*x*_eff_/d*t* = 0, thus allowing us to write the fixed points of Eq. **4** besides *x*_eff_ = 0 as

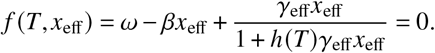

We rewrite this as

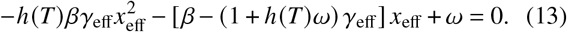

Eq. **13** has a quadratic term with a negative leading coefficient, indicating a downward-opening parabola. For the system to simultaneously exhibit an unstable equilibrium 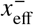 (where 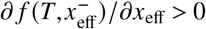) and a stable equilibrium 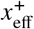 (where 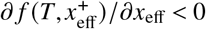), the discriminant Δ must be greater than 0, where Δ is defined by Eq. **13** as

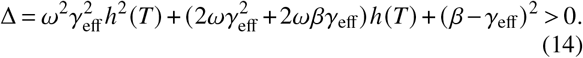

We fix all other parameters and only vary the temperature *T* . It can be shown that when 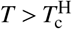, the discriminant Δ is greater than 0, where

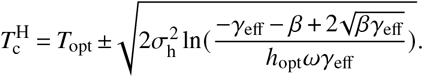

In this case, all species eventually go extinct due to the extreme temperature at this point.

Generally, whether an ecosystem will face collapse also depends on the species initial abundances. We denote *T*^L^ as the tipping temperature marking the transition from low initial abundances to a loss of resilience. This is defined as the maximum temperature for which 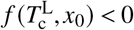 holds consistently, as derived from Eq. **13**,

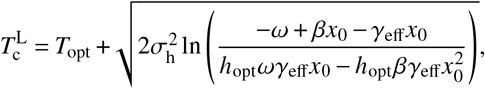

where 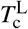 defines a tipping point at the low initial abundances of species. We next define the low initial abundances. If the initial abundances fall below the minimum of the unstable fixed point, the species will inevitably enter the extinction basin of attraction, which implies that the lower bound *a* of low initial abundances is 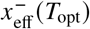 where 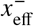 is the unstable fixed point, namely,

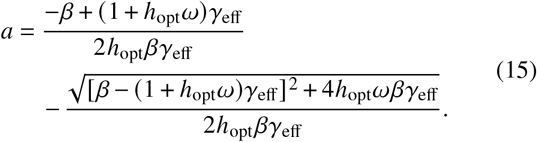

Conversely, if the initial abundances exceed the maximum of the unstable fixed point, species enter the survival basin of attraction, and the system’s critical behavior becomes independent of initial conditions. This indicates that the upper bound of low initial abundances corresponds to the point *b* at which ∂ *f* (*T, b*)/∂*x*_eff_ = 0, so

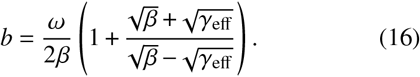

Thus, low initial abundances are defined within [*a, b*].

### Basin of attraction in the 1D model

We focus on the case Δ > 0, as this regime yields a basin of attraction for the system. Since temperature affects the system only through the handling times *h*, we first examine the direct impact of *h* for simplicity. From Eq. **13**, the unstable fixed point of the system is

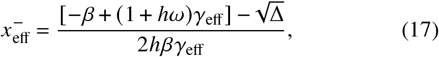

and the basin of attraction for survival is given by 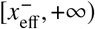.

Next, we differentiate 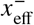 with respect to *h*, yielding

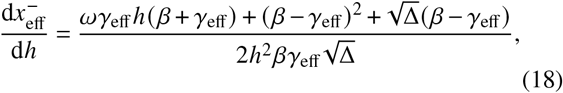

where Δ defined as in Eq. **14**, can be simplified to 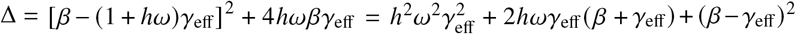. To determine the sign of Eq. **18**, it suffices to examine the sign of its numerator *z*. We then substitute the first term (*ωγ*_eff_ *h*(*β* + *γ*_eff_)) with an expression containing Δ, resulting in

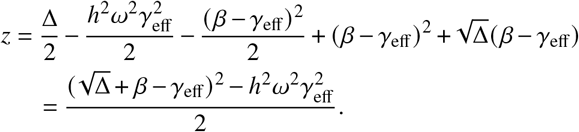

Since *γ*_eff_ ≫ *β*, we obtain 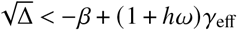. It follows that 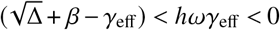, so 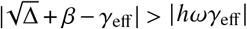, and hence 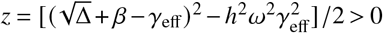. Thus, 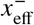 increases with *h*, indicating that faster warming prolongs handling times and shrinks the survival basin 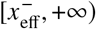. This implies that if initial abundances fail to track shifts in the basin, extinction risk increases.

## Supporting information

SOM

## Data availability

All ecological network data supporting the findings of this study are publicly available from the Web of Life database at https://www.web-of-life.es/. The Shared Socioeconomic Pathways (SSPs) data are accessible through the Earth System Grid Federation (ESGF) CMIP6 search portal at https://esgf-node.llnl.gov/search/cmip6/.

## Code availability

The simulation code supporting this work is available for download at https://doi.org/10.5281/zenodo.18437969.

## Acknowledgements

This work was supported by the National Natural Science Foundation of China (Grants No. 12271519), the Fundamental Research Funds for the Central Universities (Grant Nos. 2024KYJD2002 and 2023ZDYQ11005) and the NERC Pushing the Frontiers (CNE/X013766/1). For the purpose of open access, the author has applied a CC BY public copyright licence to any author accepted manuscript arising from this submission.

