## Supplementary material for "Tipping Points and Emergent Phases of Species Diversity in Mutualistic Ecological Networks under Global Warming": SOM

#### 1 Saturating mutualistic functional response

Since Holling's seminal "disk experiment" in 1959 introduced the concept of handling time $h$, his functional response model has provided a quantitative framework for understanding predator–prey interactions. Handling time–the period during which a predator spends capturing, consuming, and digesting prey before resuming hunting–underpins the saturating form of Type II functional responses [1]. This concept was later extended to mutualistic systems to include handling time associated with biological processes such as resource processing and pollination [2, 3]. In mutualistic systems, partners exchange resources or services, and the benefits derived from mutualists typically saturate with increasing partner density or investment, giving rise to the "saturating mutualistic functional response" model. Recent work shows that temperature fluctuations can strongly affect handling time in biological processes [4, 5], prompting growing interest in how temperature modulates mutualistic interactions. Modeling handling time as a temperature–dependent parameter offers a more realistic representation of how environmental variation shapes the strength of mutualism.

To explore the effects of global warming, we introduce temperature-dependent saturation into the mutualistic network model described in the main text

$$\frac{\mathrm{d}P_i}{\mathrm{d}t} = P_i\left[\omega_i^{(P)} - \sum_{j=1}^{S_P}\beta_{ij}^{(P)}P_j + \frac{\sum_{k=1}^{S_A}\gamma_{ik}^{(P)}A_k}{1+h(T)\sum_{k=1}^{S_A}\gamma_{ik}^{(P)}A_k}\right],$$
$$\frac{\mathrm{d}A_i}{\mathrm{d}t} = A_i\left[\omega_i^{(A)} - \sum_{j=1}^{S_A}\beta_{ij}^{(A)}A_j + \frac{\sum_{k=1}^{S_P}\gamma_{ik}^{(A)}P_k}{1+h(T)\sum_{k=1}^{S_P}\gamma_{ik}^{(A)}P_k}\right]. \tag{A1}$$

An equivalent form of the last term on the right-hand side of Eq. A1 is

$$M = \frac{P}{1+h(T)P}, \tag{A2}$$

where $M$ represents mutualistic benefit, $P$ denotes partner density, and the function $h(T)$ quantifies the handling time, reflecting a species' capacity to utilize the benefits provided by its partners. Specifically,

$$h(T) = h_{\mathrm{opt}}\exp\left(\frac{(T-T_{\mathrm{opt}})^2}{2\sigma_{\mathrm{h}}^2}\right), \tag{A3}$$

where $h_{\mathrm{opt}}$ denotes the value of $h(T)$ at the optimal temperature $T_{\mathrm{opt}}$, and $\sigma_{\mathrm{h}}$ denotes the species' thermal tolerance range.

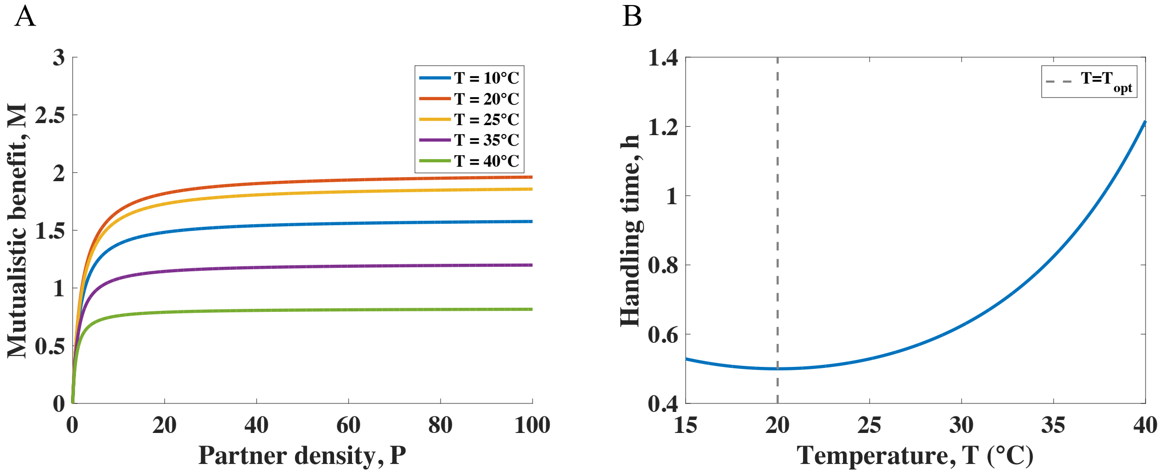


Figure S1: Global warming increases handling time, ultimately leading to a saturating decline in mutualistic interactions. (*A*) The relationship between partner density and mutualistic benefit shows a saturating mutualistic response across temperatures: at low partner density, the mutualistic benefit increases approximately linearly, whereas at high partner density, the species becomes "service-saturated". (*B*) Handling time varies unimodally with temperature. The greater the deviation from the optimal temperature, the longer the handing time required. Here, $h_{\mathrm{opt}} = 0.5$, $\sigma_{\mathrm{h}} = 15$.

Figure S1*A* shows the relationship between partner density and mutualistic benefit across temperatures. Benefit rises linearly at low densities but saturates at high densities. Temperature modulates the handling time $h$, which captures a species' capacity to process benefits from its mutualistic partners. To explore how $h(T)$ varies with temperature, Fig. S1*B* depicts the dependence of handling time on environmental temperature. The handling time $h(T)$ is minimal at the optimal temperature $T_{\text{opt}}$ and increases symmetrically as temperature deviates from this optimum. This prolongation of handling time accelerates saturation and ultimately reduces the potential mutualistic benefit under thermal stress.

### 2 Mean-field theory

To gain deeper insights into tipping points, we perform a rigorous theoretical analysis. For model A1, isolating intraspecific competition, it can be rewritten as

$$\begin{aligned}\frac{\mathrm{d}P_i}{\mathrm{d}t} &= P_i\left[\omega_i^{(P)} - \beta_{ii}^{(P)}P_i - \sum_{j\neq i}\beta_{ij}^{(P)}P_j + \frac{\sum_k \gamma_{ik}^{(P)}A_k}{1+h(T)\sum_k \gamma_{ik}^{(P)}A_k}\right],\\ \frac{\mathrm{d}A_i}{\mathrm{d}t} &= A_i\left[\omega_i^{(A)} - \beta_{ii}^{(A)}A_i - \sum_{j\neq i}\beta_{ij}^{(A)}A_j + \frac{\sum_k \gamma_{ik}^{(A)}P_k}{1+h(T)\sum_k \gamma_{ik}^{(A)}P_k}\right].\end{aligned} \tag{A4}$$

We will assume that the self-growth rate and intraspecific competition are the same for all species [6]. Namely, $\omega_i^{(P)} = \omega_i^{(A)} = \omega$, $\beta_{ii}^{(P)} = \beta_{ii}^{(A)} = \beta_0$. Hence, we obtain

$$\begin{aligned}\frac{\mathrm{d}P_i}{\mathrm{d}t} &= P_i\left[\omega - \beta_0 P_i - \sum_{j\neq i}\beta_{ij}^{(P)}P_j + \frac{\sum_k \gamma_{ik}^{(P)}A_k}{1+h(T)\sum_k \gamma_{ik}^{(P)}A_k}\right],\\ \frac{\mathrm{d}A_i}{\mathrm{d}t} &= A_i\left[\omega - \beta_0 A_i - \sum_{j\neq i}\beta_{ij}^{(A)}A_j + \frac{\sum_k \gamma_{ik}^{(A)}P_k}{1+h(T)\sum_k \gamma_{ik}^{(A)}P_k}\right].\end{aligned} \tag{A5}$$

From the above formula, we observe that node activity is governed by nearest neighbors through the interaction term, so we focus on quantities properties of the average nearest neighbor. We treat other nodes' impact on $i$ as a scalar $y_i$, with nearest-neighbor mean $\langle y_i\rangle_{nn}$. Setting $y_j(x_i) = U(x_i, x_j)$ represents that for neighbor $j$, its influence on $i$ is $y_j(x_i)$; using the adjacency matrix $A$, the total neighbor influence on $i$ is $\sum_{j\neq i} A_{ij}U(x_i, x_j) = s_i\langle y_i\rangle_{nn}$ where $s_i = \sum_{j\neq i} A_{ij}$. To facilitate analysis, we introduce an averaging procedure [7]: for node $i$, neighbor $j$ is selected with probability proportional to $j$'s connectivity, nodes with larger degree are more likely to be chosen so that their weight in $\langle y_i\rangle_{nn}$ is proportional to $s_j$, so we introduce the operator $\langle y_i\rangle_{nn} = \mathcal{L}(M, \mathbf{y}) = \mathbf{1}^\top M\mathbf{y}/\mathbf{1}^\top M\mathbf{1}$ as the influence from the average nearest neighbor, where $M = (M_{ij})$ is the network interaction matrix. Additionally, we define $s_i^{\beta^{(P)}} = \sum_{j\neq i}\beta_{ij}^{(P)}$, $s_i^{\beta^{(A)}} = \sum_{j\neq i}\beta_{ij}^{(A)}$, $s_i^{\gamma^{(P)}} = \sum_k \gamma_{ik}^{(P)}$, $s_i^{\gamma^{(A)}} = \sum_k \gamma_{ik}^{(A)}$. We then use this operator $\mathcal{L}(M, \mathbf{y})$ to write Eq. A5 as

$$\begin{aligned}\frac{\mathrm{d}P_i}{\mathrm{d}t} &= P_i\left[\omega - \beta_0 P_i - s_i^{\beta^{(\mathrm{P})}}\mathcal{L}(\beta^{(P)}, P) + \frac{s_i^{\gamma^{(P)}}\mathcal{L}(\gamma^{(P)}, A)}{1+h(T)s_i^{\gamma^{(P)}}\mathcal{L}(\gamma^{(P)}, A)}\right],\\ \frac{\mathrm{d}A_i}{\mathrm{d}t} &= A_i\left[\omega - \beta_0 A_i - s_i^{\beta^{(A)}}\mathcal{L}(\beta^{(A)}, A) + \frac{s_i^{\gamma^{(A)}}\mathcal{L}(\gamma^{(A)}, P)}{1+h(T)s_i^{\gamma^{(A)}}\mathcal{L}(\gamma^{(A)}, P)}\right].\end{aligned} \tag{A6}$$

With four weight matrices $\beta^{(P)}$, $\beta^{(A)}$, $\gamma^{(P)}$, $\gamma^{(A)}$ as inputs to the mean-field operator in Eq. A6, the high-dimensional dynamics in Eq. A1 can be condensed into a reduced four-dimensional system

$$
\begin{aligned}
\frac{\mathrm{d}\mathcal{L}(\beta^{(P)},P)}{\mathrm{d}t} &\approx \mathcal{L}(\beta^{(P)},P)\bigg[\omega-\beta_0\mathcal{L}(\beta^{(P)},P)-\mathcal{L}(\beta^{(P)},\mathbf{s}^{\beta^{(\mathbf{P})}})\\
&\qquad \mathcal{L}(\beta^{(P)},P)+\frac{\mathcal{L}(\beta^{(P)},\mathbf{s}^{\gamma^{(\mathbf{P})}})\mathcal{L}(\gamma^{(P)},A)}{1+h(T)\mathcal{L}(\beta^{(P)},\mathbf{s}^{\gamma^{(\mathbf{P})}})\mathcal{L}(\gamma^{(P)},A)}\bigg],\\
\frac{\mathrm{d}\mathcal{L}(\beta^{(A)},A)}{\mathrm{d}t} &\approx \mathcal{L}(\beta^{(A)},A)\bigg[\omega-\beta_0\mathcal{L}(\beta^{(A)},A)-\mathcal{L}(\beta^{(A)},\mathbf{s}^{\beta^{(\mathbf{A})}})\\
&\qquad \mathcal{L}(\beta^{(A)},A)+\frac{\mathcal{L}(\beta^{(A)},\mathbf{s}^{\gamma^{(\mathbf{A})}})\mathcal{L}(\gamma^{(A)},P)}{1+h(T)\mathcal{L}(\beta^{(A)},\mathbf{s}^{\gamma^{(\mathbf{A})}})\mathcal{L}(\gamma^{(A)},P)}\bigg],\\
\frac{\mathrm{d}\mathcal{L}(\gamma^{(A)},P)}{\mathrm{d}t} &\approx \mathcal{L}(\gamma^{(A)},P)\bigg[\omega-\beta_0\mathcal{L}(\gamma^{(A)},P)-\mathcal{L}(\gamma^{(A)},\mathbf{s}^{\beta^{(\mathbf{P})}})\\
&\qquad \mathcal{L}(\beta^{(P)},P)+\frac{\mathcal{L}(\gamma^{(A)},\mathbf{s}^{\gamma^{(\mathbf{P})}})\mathcal{L}(\gamma^{(P)},A)}{1+h(T)\mathcal{L}(\gamma^{(A)},\mathbf{s}^{\gamma^{(\mathbf{P})}})\mathcal{L}(\gamma^{(P)},A)}\bigg],\\
\frac{\mathrm{d}\mathcal{L}(\gamma^{(P)},A)}{\mathrm{d}t} &\approx \mathcal{L}(\gamma^{(P)},A)\bigg[\omega-\beta_0\mathcal{L}(\gamma^{(P)},A)-\mathcal{L}(\gamma^{(P)},\mathbf{s}^{\beta^{(\mathbf{A})}})\\
&\qquad \mathcal{L}(\beta^{(A)},A)+\frac{\mathcal{L}(\gamma^{(P)},\mathbf{s}^{\gamma^{(\mathbf{A})}})\mathcal{L}(\gamma^{(A)},P)}{1+h(T)\mathcal{L}(\gamma^{(P)},\mathbf{s}^{\gamma^{(\mathbf{A})}})\mathcal{L}(\gamma^{(A)},P)}\bigg].
\end{aligned}
\tag{A7}
$$

In order to simplify the calculation notation, let $x_1=\mathcal{L}(\beta^{(P)},P), x_2=\mathcal{L}(\beta^{(A)},A), x_3=\mathcal{L}(\gamma^{(A)},P), x_4=\mathcal{L}(\gamma^{(P)},A)$. Here, $x_1$ and $x_2$ represent the weighted effects of interspecific competition on the abundances of plants and pollinators, respectively; $x_3$ and $x_4$ denote the weighted average effects of pollinators and plants on plant abundance $P$ and pollinator abundance $A$ through mutualistic interactions, respectively. Similarly, we set $\beta_1=\mathcal{L}(\beta^{(P)},\mathbf{s}^{\beta^{(\mathbf{P})}}), \beta_2=\mathcal{L}(\beta^{(A)},\mathbf{s}^{\beta^{(\mathbf{A})}}), \beta_3=\mathcal{L}(\gamma^{(A)},\mathbf{s}^{\beta^{(\mathbf{P})}}), \beta_4=\mathcal{L}(\gamma^{(P)},\mathbf{s}^{\beta^{(\mathbf{A})}}), \gamma_1=\mathcal{L}(\beta^{(P)},\mathbf{s}^{\gamma^{(\mathbf{P})}}), \gamma_2=\mathcal{L}(\beta^{(A)},\mathbf{s}^{\gamma^{(\mathbf{A})}}), \gamma_3=\mathcal{L}(\gamma^{(A)},\mathbf{s}^{\gamma^{(\mathbf{P})}}), \gamma_4=\mathcal{L}(\gamma^{(P)},\mathbf{s}^{\gamma^{(\mathbf{A})}})$. Then

$$
\begin{aligned}
\frac{\mathrm{d}x_1}{\mathrm{d}t} &= x_1\left(\omega-\beta_0x_1-\beta_1x_1+\frac{\gamma_1x_4}{1+h(T)\gamma_1x_4}\right),\\
\frac{\mathrm{d}x_2}{\mathrm{d}t} &= x_2\left(\omega-\beta_0x_2-\beta_2x_2+\frac{\gamma_2x_3}{1+h(T)\gamma_2x_3}\right),\\
\frac{\mathrm{d}x_3}{\mathrm{d}t} &= x_3\left(\omega-\beta_0x_3-\beta_3x_1+\frac{\gamma_3x_4}{1+h(T)\gamma_3x_4}\right),\\
\frac{\mathrm{d}x_4}{\mathrm{d}t} &= x_4\left(\omega-\beta_0x_4-\beta_4x_2+\frac{\gamma_4x_3}{1+h(T)\gamma_4x_3}\right).
\end{aligned}
\tag{A8}
$$

Following the analysis above, the high-dimensional space can be reduced to four dimensions. Expressing Eq. A8 in vector form yields

$$
\frac{\mathrm{d}\mathbf{x}}{\mathrm{d}t}=\mathbf{x}\odot(\omega-\beta_0\mathbf{x}-B\mathbf{x}+G\mathbf{x}\oslash(1+h(T)G\mathbf{x})),
\tag{A9}
$$

where $\mathbf{x}=(x_1,x_2,x_3,x_4)^T$, $\odot$ and $\oslash$ respectively denote element-wise multiplication and element-wise division, and the specific forms of matrices $B$ and $G$ are as follows

$$
B=\begin{bmatrix}\beta_1 & 0 & 0 & 0\\ 0 & \beta_2 & 0 & 0\\ \beta_3 & 0 & 0 & 0\\ 0 & \beta_4 & 0 & 0\end{bmatrix},\qquad G=\begin{bmatrix}0 & 0 & 0 & \gamma_1\\ 0 & 0 & \gamma_2 & 0\\ 0 & 0 & 0 & \gamma_3\\ 0 & 0 & \gamma_4 & 0\end{bmatrix}.
\tag{A10}
$$

We will focus on the influence of the nearest neighbors, which simplifies this system to

$$\frac{\mathrm{d}\mathbf{x}}{\mathrm{d}t} = \mathbf{x} \odot \left(\omega - \beta_0 \mathbf{x} - \mathbf{b}\mathcal{L}(Z, \mathbf{x}) + \mathbf{g}\mathcal{L}(Z, \mathbf{x}) \oslash (1 + h(T)\mathbf{g}\mathcal{L}(Z, \mathbf{x}))\right), \tag{A11}$$

where

$$Z = \begin{bmatrix} \beta_1 & 0 & 0 & \gamma_1 \\ 0 & \beta_2 & \gamma_2 & 0 \\ \beta_3 & 0 & 0 & \gamma_3 \\ 0 & \beta_4 & \gamma_4 & 0 \end{bmatrix}, \quad \mathbf{b} = \begin{bmatrix} \beta_1 \\ \beta_2 \\ \beta_3 \\ \beta_4 \end{bmatrix}, \quad \mathbf{g} = \begin{bmatrix} \gamma_1 \\ \gamma_2 \\ \gamma_3 \\ \gamma_4 \end{bmatrix}. \tag{A12}$$

Here, $Z$ denotes the adjacency matrix formed by reconstructing the four-dimensional information derived by Eq. A8. Similarly, we can replace $M$ by $Z$ as the input matrix of the operator $\mathcal{L}(M, \mathbf{y})$ to Eq. A11. Finally, we obtain

$$\frac{\mathrm{d}x_{\text{eff}}}{\mathrm{d}t} = x_{\text{eff}} \left(\omega - \beta x_{\text{eff}} + \frac{\gamma_{\text{eff}} x_{\text{eff}}}{1 + h(T)\gamma_{\text{eff}} x_{\text{eff}}}\right), \tag{A13}$$

where $\beta = \beta_{\text{eff}} + \beta_0$, and

$$\begin{aligned} x_{\text{eff}} &= \frac{\mathbf{1}^\top Z \left[x_1, x_2, x_3, x_4\right]^\top}{\mathbf{1}^\top Z \mathbf{1}}, \\ \beta_{\text{eff}} &= \frac{\mathbf{1}^\top Z \left[\beta_1, \beta_2, \beta_3, \beta_4\right]^\top}{\mathbf{1}^\top Z \mathbf{1}}, \\ \gamma_{\text{eff}} &= \frac{\mathbf{1}^\top Z \left[\gamma_1, \gamma_2, \gamma_3, \gamma_4\right]^\top}{\mathbf{1}^\top Z \mathbf{1}}. \end{aligned} \tag{A14}$$

Here, $x_{\text{eff}}$ denotes the effective average abundance of plants and pollinators, as viewed from their shared perspective. Similarly, $\beta = \beta_{\text{eff}} + \beta_0$ characterizes the effective average competition, combining interspecific and intraspecific effects, with $\beta_{\text{eff}}$ representing effective average interspecific competition. The parameters $\omega$ and $\gamma_{\text{eff}}$ correspond to the effective average intrinsic growth rate and mutualistic strength, respectively. Consequently, the network of interspecific competition and mutualistic interactions in the microscopic description can be collapsed into the two macroscopic resilience parameters $\beta_{\text{eff}}$ and $\gamma_{\text{eff}}$. This reduction allows the bifurcation analysis framework–originally developed for low-dimensional systems–to be applied to high-dimensional mutualistic networks under temperature influence, enabling accurate prediction of system responses to perturbations and precise identification of tipping points at which resilience is lost.

### 3 Calculation of tipping points

To identify the tipping points leading to species extinction, we perform bifurcation analysis on Eq. A13. At steady state ($\mathrm{d}x_{\text{eff}}/\mathrm{d}t = 0$), the equilibrium points of Eq. A13, excluding $x_{\text{eff}} = 0$, are given by the function

$$f(T, x_{\text{eff}}) = \omega - \beta x_{\text{eff}} + \frac{\gamma_{\text{eff}} x_{\text{eff}}}{1 + h(T)\gamma_{\text{eff}} x_{\text{eff}}} = 0. \tag{A15}$$

Hence we have the quadratic equation

$$-h(T)\beta\gamma_{\text{eff}} x_{\text{eff}}^2 - \left[\beta - (1 + h(T)\omega)\gamma_{\text{eff}}\right] x_{\text{eff}} + \omega = 0, \tag{A16}$$

which has a negative leading coefficient (indicating a downward-opening parabola). Its equilibrium points are

$$x_{\text{eff}}^{\pm}(T) = \frac{[-\beta + (1 + h(T)\omega)\gamma_{\text{eff}}]}{2h(T)\beta\gamma_{\text{eff}}} \pm \frac{\sqrt{\Delta}}{2h(T)\beta\gamma_{\text{eff}}}, \tag{A17}$$

where $\Delta = [\beta - (1 + h(T)\omega)\gamma_{\text{eff}}]^2 + 4h(T)\omega\beta\gamma_{\text{eff}}$.

To assess the resilience of the equilibrium points, we first compute the derivative of the function with respect to $x_{\text{eff}}$

$$\partial f(T, x_{\text{eff}})/\partial x_{\text{eff}} = -\beta + \frac{\gamma_{\text{eff}}}{(1 + h(T)\gamma_{\text{eff}}x_{\text{eff}})^2}. \tag{A18}$$

Substituting $x_{\text{eff}}^{-}$ into the above equation yields

$$\begin{aligned}
\partial f(T, x_{\text{eff}}^{-})/\partial x_{\text{eff}} &= -\beta + \frac{\gamma_{\text{eff}}}{(1 + h(T)\gamma_{\text{eff}}x_{\text{eff}}^{-})^2} \\
&= -\beta + \frac{\gamma_{\text{eff}}}{(1 + h(T)\gamma_{\text{eff}} \cdot \frac{[-\beta + (1+h(T)\omega)\gamma_{\text{eff}}] - \sqrt{\Delta}}{2h(T)\beta\gamma_{\text{eff}}})^2} \\
&= -\beta + \frac{\gamma_{\text{eff}}}{(1 + \frac{[-\beta + (1+h(T)\omega)\gamma_{\text{eff}}] - \sqrt{\Delta}}{2\beta})^2} \\
&= -\beta + \frac{\gamma_{\text{eff}}}{(\frac{[\beta + (1+h(T)\omega)\gamma_{\text{eff}}] - \sqrt{\Delta}}{2\beta})^2} \\
&= -\beta + \frac{4\gamma_{\text{eff}}\beta^2}{([\beta + (1 + h(T)\omega)\gamma_{\text{eff}}] - \sqrt{\Delta}})^2 \\
&= \frac{\beta}{([\beta + (1 + h(T)\omega)\gamma_{\text{eff}}] - \sqrt{\Delta})^2} \\
&\cdot (-([\beta + (1 + h(T)\omega)\gamma_{\text{eff}}] - \sqrt{\Delta})^2 + 4\gamma_{\text{eff}}\beta).
\end{aligned} \tag{A19}$$

From the above equation, the sign of $\partial f(T, x_{\text{eff}}^{-})/\partial x_{\text{eff}}$ is determined solely by the second term

$$\begin{aligned}
&-([\beta + (1 + h(T)\omega)\gamma_{\text{eff}}] - \sqrt{\Delta})^2 + 4\gamma_{\text{eff}}\beta \\
&= 2[-\beta^2 - \gamma_{\text{eff}}^2 - 2h(T)\omega\gamma_{\text{eff}}^2 - h(T)^2\omega^2\gamma_{\text{eff}}^2 \\
&\quad +2\beta\gamma_{\text{eff}} - 2h(T)\omega\gamma_{\text{eff}}\beta + [\beta + (1 + h(T)\omega)\gamma_{\text{eff}}]\sqrt{\Delta}] \\
&= 2[-\Delta + [\beta + (1 + h(T)\omega)\gamma_{\text{eff}}]\sqrt{\Delta}] \\
&= 2[\sqrt{\Delta}[-\sqrt{\Delta} + \beta + (1 + h(T)\omega)\gamma_{\text{eff}}]].
\end{aligned} \tag{A20}$$

Since $\gamma_{\text{eff}} \gg \beta$, we obtain $\sqrt{\Delta} < -\beta + (1 + h\omega)\gamma_{\text{eff}}$. So it follows that

$$\begin{aligned}
&2\sqrt{\Delta}\big[-\sqrt{\Delta} + \beta + (1 + h(T)\omega)\gamma_{\text{eff}}\big] \\
&\qquad > 2\sqrt{\Delta}\big[\beta - (1 + h(T)\omega)\gamma_{\text{eff}} + \beta + (1 + h(T)\omega)\gamma_{\text{eff}}\big] \\
&\qquad = 4\beta\sqrt{\Delta} > 0,
\end{aligned} \tag{A21}$$

so $\partial f(T, x_{\text{eff}}^{-})/\partial x_{\text{eff}} > 0$. Thus, $x_{\text{eff}}^{-}$ corresponds to the unstable point. Similarly, evaluating

$\partial f(T, x_{\text{eff}}^{+})/\partial x_{\text{eff}} < 0$ gives

$$\begin{aligned}\partial f(T, x_{\text{eff}}^{+})/\partial x_{\text{eff}} &= -\beta + \frac{\gamma_{\text{eff}}}{(1 + h(T)\gamma_{\text{eff}} x_{\text{eff}}^{+})^2} \\ &= \frac{\beta}{([\beta + (1 + h(T)\omega)\gamma_{\text{eff}}] + \sqrt{\Delta})^2} \\ &\cdot (-([\beta + (1 + h(T)\omega)\gamma_{\text{eff}}] + \sqrt{\Delta})^2 + 4\gamma_{\text{eff}}\beta) \\ &= \frac{2\beta}{([\beta + (1 + h(T)\omega)\gamma_{\text{eff}}] + \sqrt{\Delta})^2} \\ &\cdot (-\Delta - (\beta + (1 + h(T)\omega)\gamma_{\text{eff}})\sqrt{\Delta}) < 0,\end{aligned} \tag{A22}$$

so, $\partial f(T, x_{\text{eff}}^{+})/\partial x_{\text{eff}} < 0$. In this case, the system is resilient. Therefore, as shown in Fig. 5*A* of the main text, for the system to exhibit both an unstable equilibrium $x_{\text{eff}}^{-}$ (where $f(T, x_{\text{eff}}^{-}) = 0$ and $\partial f(T, x_{\text{eff}}^{-})/\partial x_{\text{eff}} > 0$) and a stable equilibrium $x_{\text{eff}}^{+}$ (where $f(T, x_{\text{eff}}^{+}) = 0$ and $\partial f(T, x_{\text{eff}}^{+})/\partial x_{\text{eff}} < 0$), the discriminant $\Delta$ must be positive (otherwise, the system will not have any non-trivial real equilibrium point), where $\Delta$ is defined by Eq. A16 as

$$\Delta = \omega^2\gamma_{\text{eff}}^2 h^2(T) + (2\omega\gamma_{\text{eff}}^2 + 2\omega\beta\gamma_{\text{eff}})h(T) + (\beta - \gamma_{\text{eff}})^2. \tag{A23}$$

It is known that $h(T)$ increases monotonically with $T$ for $T > T_{\text{opt}}$. Therefore, we first calculate the critical value of $h(T)$ for which $\Delta = 0$, namely

$$h(T) = \frac{-\gamma_{\text{eff}} - \beta \pm 2\sqrt{\beta\gamma_{\text{eff}}}}{\omega\gamma_{\text{eff}}}. \tag{A24}$$

From Fig. 5 *A* in the main text, we find that when $h(T)$ (which increases with $T$ for $T > T_{\text{opt}}$) exceeds a ctitical value, $f(T, x_{\text{eff}}) < 0$ for all conditions and all species eventually go extinct as temperature pressure becomes overwhelming (0 is the only fixed point of Eq. A13). Since this critical value corresponds to the maximum handling time that prevents species extinction, and given that $\gamma_{\text{eff}} \gg \beta$, $0 < h_{\text{opt}} < 1$, and $-1 \leq \omega < 0$, we substitute $h(T) = h_{\text{opt}} \exp\left((T - T_{\text{opt}})^2/2\sigma_{\text{h}}^2\right)$ into Eq. A24, yielding

$$T_{\text{c}}^{\text{H}} = T_{\text{opt}} + \sqrt{2\sigma_{\text{h}}^2 \ln\left(\frac{-\gamma_{\text{eff}} - \beta + 2\sqrt{\beta\gamma_{\text{eff}}}}{h_{\text{opt}}\omega\gamma_{\text{eff}}}\right)}. \tag{A25}$$

After this tipping point, all species ultimately go extinct due to the extreme temperature.

Generally, the risk of ecosystem collapse depends not only on environmental temperature but also on species' initial abundances. As shown in Fig. 5*A* of the main text, when initial abundances are low, the system exhibits reduced resilience and slower recovery from disturbances. Consequently, different initial states correspond to different temperature thresholds for avoiding collapse. For a fixed initial abundance, there exists a specific temperature: the system gradient is positive when below this temperature, but turns negative once it is exceeded. We denote $T_{\text{c}}^{\text{L}}$ as the tipping temperature marking the transition from low initial abundances to a loss of resilience. This is defined as the maximum temperature for which $f(T_{\text{c}}^{\text{L}}, x_0 < 0$ holds consistently, as derived from Eq. A15

$$T_{\text{c}}^{\text{L}} = T_{\text{opt}} + \sqrt{2\sigma_{\text{h}}^2 \ln\left(\frac{-\omega + \beta x_0 - \gamma_{\text{eff}} x_0}{h_{\text{opt}}\omega\gamma_{\text{eff}} x_0 - h_{\text{opt}}\beta\gamma_{\text{eff}} x_0^2}\right)}. \tag{A26}$$

Next, we derive the interval corresponding to low initial abundances. The unstable equilibrium point is given by

$$x_{\text{eff}}^{-}(T) = \frac{[-\beta + (1 + h(T)\omega)\gamma_{\text{eff}}]}{2h(T)\beta\gamma_{\text{eff}}} - \frac{\sqrt{\Delta}}{2h(T)\beta\gamma_{\text{eff}}}. \tag{A27}$$

If the initial abundances fall below the unstable equilibrium minimum, the species will inevitably enter the extinction basin of attraction. This implies that the left endpoint of the low-abundance interval is $x_{\text{eff}}^{-}(T_{\text{opt}})$, namely,

$$a = \frac{[-\beta + (1 + h_{\text{opt}}\omega)\gamma_{\text{eff}}]}{2h_{\text{opt}}\beta\gamma_{\text{eff}}} - \frac{\sqrt{[\beta - (1 + h_{\text{opt}}\omega)\gamma_{\text{eff}}]^2 + 4h_{\text{opt}}\omega\beta\gamma_{\text{eff}}}}{2h_{\text{opt}}\beta\gamma_{\text{eff}}}. \tag{A28}$$

Conversely, if the initial abundances exceed the unstable equilibrium maximum, the species enters the survival basin of attraction, and the system's dynamics become independent of initial abundances. This indicates that the right endpoint of the low-abundance interval is the point $b$ at which $\partial f(T, b)/\partial x_{\text{eff}} = 0$, namely,

$$b = [-\beta + (1 + h(T_{\text{c}}^{\text{H}})\omega)\gamma_{\text{eff}}]/2h(T_{\text{c}}^{\text{H}})\beta\gamma_{\text{eff}} = \frac{\omega}{2\beta}\left(1 + \frac{\sqrt{\beta} + \sqrt{\gamma_{\text{eff}}}}{\sqrt{\beta} - \sqrt{\gamma_{\text{eff}}}}\right). \tag{A29}$$

Finally, low initial abundances are defined as those in the interval $[a, b]$.

### 4 Impact of network structure on tipping points

Species interactions in nature give rise to diverse and complex network structures. Among these, Scale-Free (SF), Erdős–Rényi (ER), and Small-World (WS) networks exhibit distinct properties: SF networks follow a power-law degree distribution dominated by highly connected hubs; ER networks are randomly connected with a fixed probability; and WS networks feature high clustering and short average path lengths [7]. Bipartite networks of mutualistic plants and pollinators further display varying connection strengths. To determine which network structures are most vulnerable to global warming, we embed them within interspecific competition and mutualistic networks and then perform corresponding simulations.

To assess the influence of the interspecific competition network on warming-induced tipping points, we hold the mutualistic connectivity probability between plants and pollinators constant at $p_{\text{m}} = 0.1$. Figure S2*A* shows that, under rising temperature stress, the structure of interspecific competition networks has little effect on tipping points of resilience loss. Further, as shown in Fig. S2*B*, with both plant and pollinator networks modeled as ER networks with average degree $\langle k \rangle = 6$, tipping points under high and low initial abundances are examined as the mutualistic connectivity probability increased ($p_{\text{m}} = 0.05$ (green), 0.1 (blue), and 0.15 (red)). The results indicate that modifying mutualistic links is more effective than altering interspecific competition networks at raising temperature tipping points and delaying resilience loss, particularly in ecosystems with low initial abundances.

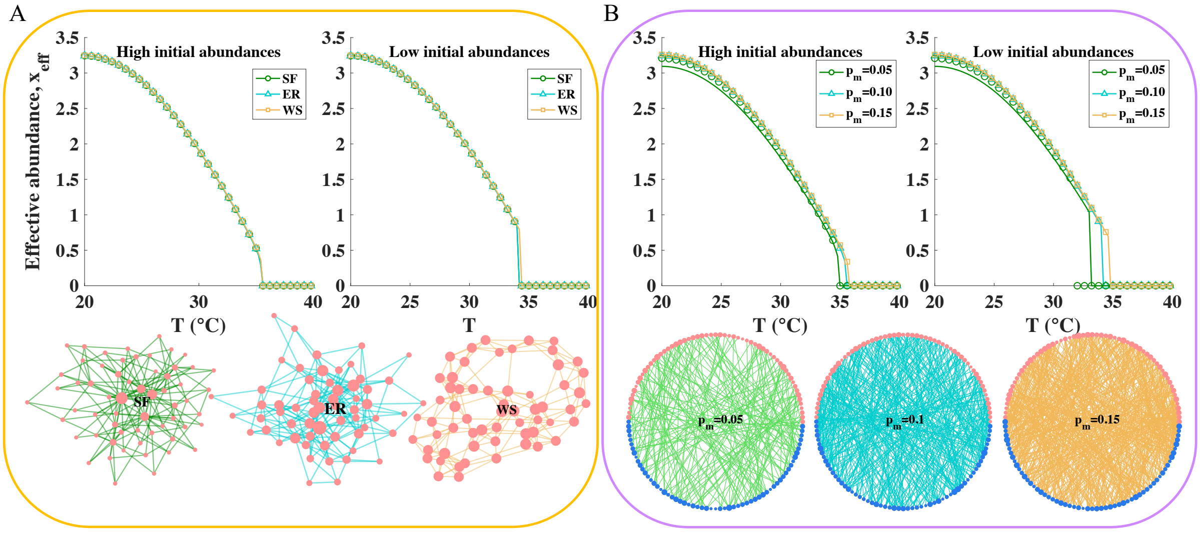


Figure S2: Temperature tipping points are only weakly influenced by interspecific competition network topology, but strongly influenced by mutualistic connectivity. ($A$) Tipping points are largely insensitive to interspecific competition network topology–SF (green), ER (blue), and WS (yellow), all with $\langle k \rangle = 6$–under high and low initial abundances, with a fixed mutualistic connectivity probability $p_{\mathrm{m}} = 0.1$. ($B$) Increasing mutualistic connectivity–$p_{\mathrm{m}} = 0.05$ (green), 0.1 (blue), and 0.15 (yellow)–raises the temperature tipping point, particularly under low initial abundances, with the interspecific competition network fixed as an ER network with average degree $\langle k \rangle = 6$. High and low initial abundances correspond to $x_0 = 2$ and $x_0 = 0.1$, respectively. Other parameters: $\beta_{ii}^{(P)} = \beta_{ii}^{(A)} = 0.3$, $\beta_{ij}^{(P)} = \beta_{ij}^{(A)} = 10^{-4}$, $\omega_i^{(P)} = \omega_i^{(A)} = -1$, $h_{\mathrm{opt}} = 0.5$, $\sigma_{\mathrm{h}} = 15$, $\tau = 0.5$, $\gamma_0 = 20$, $S_A = S_P = 60$.

### 5 Impact of variations in interaction strength

The complex behaviors of different systems under disturbance are determined by their scale, structure, and parameter space bounds [8, 9, 10]. Perturbations can alter system resilience not only by affecting environmental temperatures but also by modifying network connectivity and the strength of interactions among individuals, including intraspecific competition, interspecific competition, and mutualism. To investigate how these perturbations and temperature influence system resilience, we examine the system's dynamic response to a reduction in the fraction of mutualistic interactions $f_{\gamma_0}$ (Fig. S3$A$), an increase in the fraction of intraspecific competition interactions $f_{\beta_0}$ (Fig. S3$B$), and an increase in the fraction of interspecific competition links $f_{\mathrm{cl}}$ (Fig. S3$C$) under varying temperatures. We find that all perturbation types reduce system resilience, although resilience is higher near optimal temperatures. Among them, mutualistic interactions exert the strongest influence on system resilience.

### 6 Buffering effect of intraspecific competition on tipping points

Facing environmental stress under global warming, species typically exhibit active responses. Ecologically, this behavior is captured by describing how populations exhibit

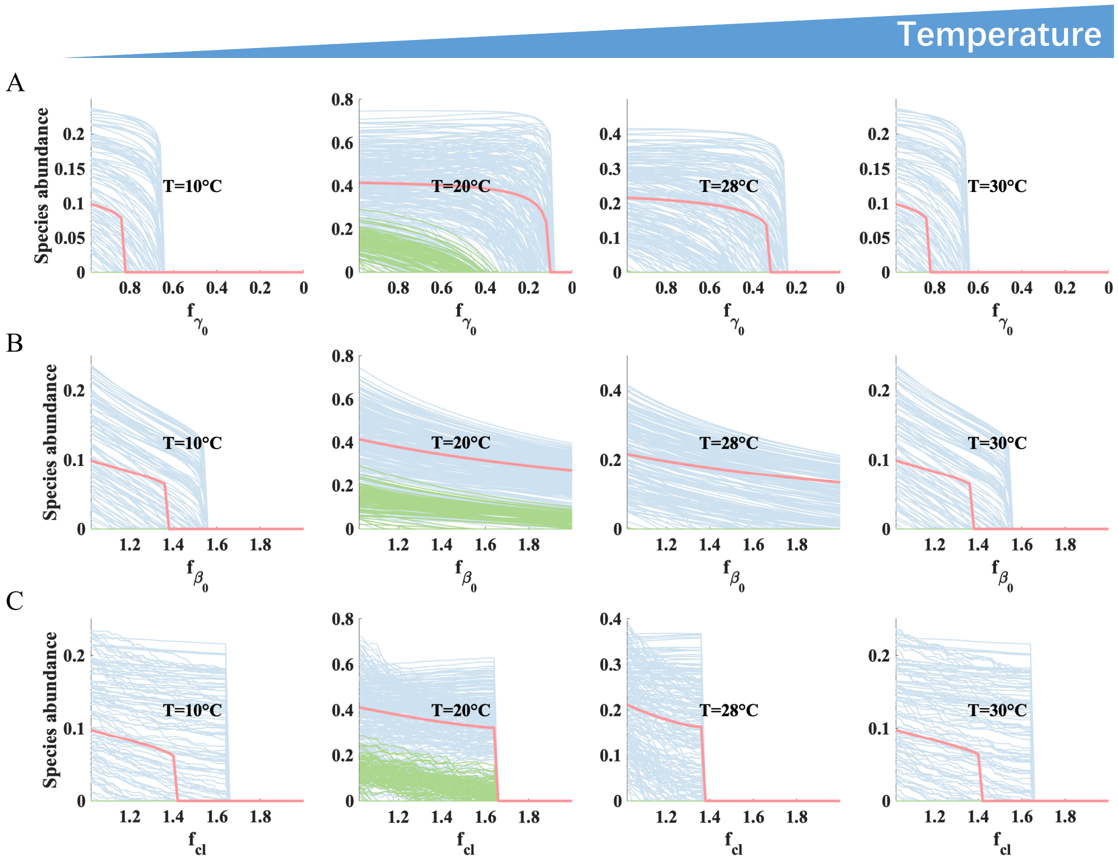


Figure S3: Reducing mutualism or increasing competition weakens system resilience, with mutualism reduction has greater influence. $(A)$ Variation in the fraction of mutualistic interactions ($f_{\gamma_0} \in [1, 0]$), $(B)$ intraspecific competition interactions ($f_{\beta_0} \in [1, 2]$), and $(C)$ interspecific competition links ($f_{\text{cl}} \in [1, 2]$) produce distinct time series dynamics of pollinators (blue), plants (green), and effective average abundances (red), with system resilience peaking near optimal temperatures. The other parameters are set as $\beta_{ii}^{(P)} = \beta_{ii}^{(A)} = 0.5$, $\beta_{ij}^{(P)} = \beta_{ij}^{(A)} = 0.01$, $\omega_i^{(P)} = \omega_i^{(A)} = -1$, $h_{\text{opt}} = 0.7$, $\sigma_{\text{h}} = 15$, $\tau = 0$, $\gamma_0 = 16$, the network is $M\_PL\_005$.

adaptive regulation in response to changing conditions, such as the Demographic Buffering Hypothesis [11] and the Stress-Gradient Hypothesis [12]. A core assumption of both hypotheses is that, under climate warming, species adaptive responses, primarily mediated through intraspecific competition, serve as a fundamental mechanism for maintaining ecosystem stability and enhancing resilience to environmental change.

Here, we assume that intraspecific competition is greatest at the temperature that optimizes reproductive performance. It reflects the fact that resource requirements peak during periods of intense reproductive effort, leading to heightened competition among individuals of the same species [13]. Under this assumption, the temperature dependence of the intraspecific competition coefficient $\beta_0(T)$ follows a unimodal response, described by a Gaussian function

$$\beta_0(T) = \beta_{\text{opt}} \exp\left(-\frac{(T - T_{\text{opt}})^2}{2s^2}\right), \tag{A30}$$

where $\beta_{\text{opt}}$ represents the maximum strength of intraspecific competition at the optimal temperature $T_{\text{opt}}$, and $s$ reflects how rapidly intraspecific competition declines as temperature deviates from the optimum. The unimodal temperature response is illustrated by the green curve in Fig. S4*A*.

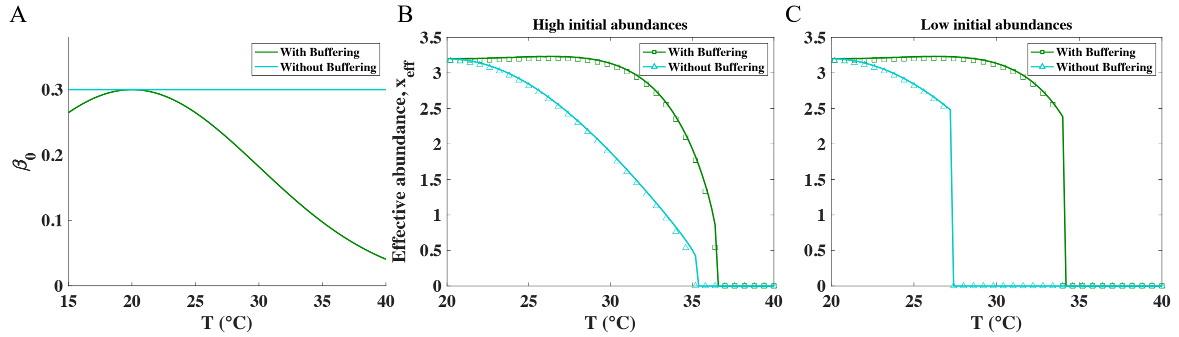


Figure S4: Adaptive intraspecific competition boosts species abundance and extends the thermal range for species coexistence to higher temperatures. (*A*) Intraspecific competition intensity ($\beta_0$) declines as temperature deviates from the optimal value. (*B*)-(*C*) Effective abundance with and without adaptive intraspecific competition at (*B*) high and (*C*) low initial abundances, showing how adaptation delays extinction. The mutualistic network structure used here is *M_PL_*005, with all other parameters as in Fig. 4 of the main text.

To investigate how ecosystems with temperature-dependent adaptive intraspecific competition respond to warming, we incorporate a unimodal thermal dependence of intraspecific competition into our model (Eq. A1) and predict the temperature thresholds at which tipping points occur under high (Fig. S4*B*) and low (Fig. S4*C*) initial abundances. The results show that, in both scenarios, species exhibit higher abundances when intraspecific competition adapts to temperature than when competition intensity is held constant. Consequently, extinction is delayed relative to the original model (Eq. A1). This adaptive response allows populations to better cope with rising temperatures, reducing the risk of species loss.

### 7 Effect of thermal tolerance on tipping points

Species' physiological tolerance varies across systems at different latitudes [14], making it essential to understand the role of latitudes in ecosystem resilience and species diversity. Variation in species' tolerance to global warming can be quantified by the thermal breadth $\sigma_{\rm h}$, as defined in Eq. A3. For example, tropical species that are adapted to long-term stable, high-temperature environments and specialized niches, tend to be more sensitive and less adaptable to warming than temperate species. Fig. S5 shows the relationship between the resilience range, $\Delta_{\rm T} = T_{\rm c} - T_{\rm opt}$, and thermal breadth $\sigma_{\rm h}$, indicating that ecosystems with broader thermal breadth tolerate warming better. These simulations validate our model and are consistent with recent findings that, despite minimal warming, tropical species face high extinction risk due to narrow thermal tolerance and near-zero thermal safety margins [15, 16]. This highlights the vulnerability of tropical species and the importance of prioritizing their conservation under climate change.

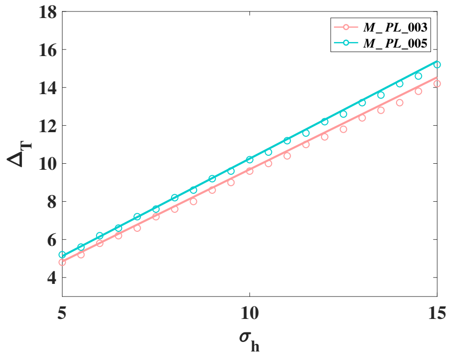


Figure S5: A proportional relationship between the resilience range ($\Delta_{\mathrm{T}} = T_{\mathrm{c}} - T_{\mathrm{opt}}$) and thermal breadth ($\sigma_{\mathrm{h}}$) indicates that broader thermal breadth (i.e., higher latitudes) enhances ecosystem resilience, as predicted theoretically (lines) and confirmed numerically (symbols). Mutualistic networks $M\_PL\_003$ and $M\_PL\_005$ are used, with all other parameters as in Fig. 2.

### 8 Basin of attraction for one-dimensional model

To understand the impact of the warming rate $\mathrm{d}T/\mathrm{d}t$ on the extinction dynamics, we consider the basin of attraction in Eq. A13. The unstable equilibrium point of the system is $x_{\mathrm{eff}}^{-}$ as in Eq. A27. It is known that $h(T)$ increases monotonically with $T$ when $T > T_{\mathrm{opt}}$. For the sake of simplicity in the analysis, we first directly examine the impact of $h$ on the system, so the unstable equilibrium point can be simplified to

$$x_{\mathrm{eff}}^{-} = \frac{[-\beta + (1+h\omega)\gamma_{\mathrm{eff}}] - \sqrt{[\beta - (1+h\omega)\gamma_{\mathrm{eff}}]^2 + 4h\omega\beta\gamma_{\mathrm{eff}}}}{2h\beta\gamma_{\mathrm{eff}}}, \tag{A31}$$

therefore, the basin of attraction for survival is $[x_{\mathrm{eff}}^{-}, +\infty]$. Next, we differentiate $x_{\mathrm{eff}}^{-}$ with respect to $h$. Set $u = [-\beta + (1+h\omega)\gamma_{\mathrm{eff}}] - \sqrt{[\beta - (1+h\omega)\gamma_{\mathrm{eff}}]^2 + 4h\omega\beta\gamma_{\mathrm{eff}}}$ and $v = 2h\beta\gamma_{\mathrm{eff}}$. Then, we have

$$\begin{aligned} \frac{\mathrm{d}v}{\mathrm{d}h} &= 2\beta\gamma_{\mathrm{eff}}, \\ \frac{\mathrm{d}u}{\mathrm{d}h} &= \omega\gamma_{\mathrm{eff}} - \frac{[\beta - (1+h\omega)\gamma_{\mathrm{eff}}] \cdot (-\omega\gamma_{\mathrm{eff}}) + 2\omega\beta\gamma_{\mathrm{eff}}}{\sqrt{[\beta - (1+h\omega)\gamma_{\mathrm{eff}}]^2 + 4h\omega\beta\gamma_{\mathrm{eff}}}}, \end{aligned} \tag{A32}$$

so

$$\begin{aligned} \frac{\mathrm{d}x_{\mathrm{eff}}^{-}}{\mathrm{d}h} &= \frac{u'v - uv'}{v^2} \\ &= \frac{\left(\omega\gamma_{\mathrm{eff}} - \frac{[\beta-(1+h\omega)\gamma_{\mathrm{eff}}]\cdot(-\omega\gamma_{\mathrm{eff}})+2\omega\beta\gamma_{\mathrm{eff}}}{\sqrt{[\beta-(1+h\omega)\gamma_{\mathrm{eff}}]^2+4h\omega\beta\gamma_{\mathrm{eff}}}}\right) \cdot 2h\beta\gamma_{\mathrm{eff}}}{(2h\beta\gamma_{\mathrm{eff}})^2} \\ &\quad - \frac{(-\beta + (1+h\omega)\gamma_{\mathrm{eff}}) \cdot 2\beta\gamma_{\mathrm{eff}}}{(2h\beta\gamma_{\mathrm{eff}})^2} \\ &\quad + \frac{\sqrt{[\beta - (1+h\omega)\gamma_{\mathrm{eff}}]^2 + 4h\omega\beta\gamma_{\mathrm{eff}}} \cdot 2\beta\gamma_{\mathrm{eff}}}{(2h\beta\gamma_{\mathrm{eff}})^2} \\ &= \frac{(\beta + \sqrt{\Delta} - \gamma_{\mathrm{eff}})\sqrt{\Delta} - \omega\gamma_{\mathrm{eff}}[\beta + (1+h\omega)\gamma_{\mathrm{eff}}]h}{2h^2\beta\gamma_{\mathrm{eff}}\sqrt{\Delta}}. \end{aligned} \tag{A33}$$

Table 1: Core characteristics of SSPs

| SSPs | Key Features | Climate Challenges |
|---|---|---|
| **SSP1** | · Gradual, global shift to sustainability<br>· Inclusive development, reduced inequality, low-carbon consumption<br>· High investments in education and health | Low challenges for mitigation (resource efficiency) and adaptation (rapid development) |
| **SSP2** | · Uneven development, environmental degradation occurs<br>· Resource intensity declines moderately<br>· No extreme socioeconomic shifts | Medium challenges to mitigation and adaptation |
| **SSP3** | · Rising nationalism, fragmented governance and security conflicts<br>· Material-intensive consumption and worsen inequality<br>· Low education and technology investment and low international priority for addressing environmental concerns | High challenges for mitigation (regionalized energy/land policies) and adaptation (slow development) |
| **SSP4** | · Extreme socioeconomic polarization<br>· High-tech economy sectors lead technology development and low-carbon investments<br>· Low-income groups remain vulnerable | Low challenges for mitigation (global high tech economy), high for adaption (regional low tech economies) |
| **SSP5** | · Market-driven, rapid economic growth and fossil fuel-abundant<br>· High tech innovation but carbon-intensive<br>· Faith in effective management of local environmental issues<br>· High investments in education and health | High challenges for mitigation (resource/fossil fuel intensive) and low for adaptation (rapid development) |

We only need to analyze the sign of the numerator in Eq. A33, namely,

$$\begin{aligned} z &= \left(\beta + \sqrt{\Delta} - \gamma_{\text{eff}}\right)\sqrt{\Delta} - \omega\gamma_{\text{eff}}\left[\beta + (1 + h\omega)\gamma_{\text{eff}}\right]h \\ &= (\beta - \gamma_{\text{eff}})\sqrt{\Delta} + \Delta - \omega\gamma_{\text{eff}}h\left[\beta + (1 + h\omega)\gamma_{\text{eff}}\right] \\ &= \omega\gamma_{\text{eff}}h(\beta + \gamma_{\text{eff}}) + (\beta - \gamma_{\text{eff}})^2 + \sqrt{\Delta}(\beta - \gamma_{\text{eff}}). \end{aligned} \tag{A34}$$

We then substitute the first term ($\omega\gamma_{\text{eff}}h(\beta + \gamma_{\text{eff}})$) with an expression containing $\Delta$, yielding

$$\begin{aligned} z &= \frac{\Delta}{2} - \frac{h^2\omega^2\gamma_{\text{eff}}^2}{2} - \frac{(\beta - \gamma_{\text{eff}})^2}{2} + (\beta - \gamma_{\text{eff}})^2 + \sqrt{\Delta}(\beta - \gamma_{\text{eff}}) \\ &= \frac{\Delta}{2} - \frac{h^2\omega^2\gamma_{\text{eff}}^2}{2} + \frac{(\beta - \gamma_{\text{eff}})^2}{2} + \sqrt{\Delta}(\beta - \gamma_{\text{eff}}) \\ &= \frac{(\sqrt{\Delta} + \beta - \gamma_{\text{eff}})^2 - h^2\omega^2\gamma_{\text{eff}}^2}{2}. \end{aligned} \tag{A35}$$

Since $\sqrt{\Delta} < -\beta + (1 + h\omega)\gamma_{\text{eff}}$, it follows that $(\sqrt{\Delta} + \beta - \gamma_{\text{eff}}) < h\omega\gamma_{\text{eff}} < 0$, so $|\sqrt{\Delta} + \beta - \gamma_{\text{eff}}| > |h\omega\gamma_{\text{eff}}|$, and hence $z = [(\sqrt{\Delta} + \beta - \gamma_{\text{eff}})^2 - h^2\omega^2\gamma_{\text{eff}}^2]/2 > 0$. Thus, $x_{\text{eff}}^-$ increases with $h$. This indicates that as the warming rate accelerates, handling times become longer, and the survival attraction basin $[x_{\text{eff}}^-, +\infty)$ determined by the unstable equilibrium point shrinks accordingly. This implies that as the initial abundances cannot keep up with this change in the survival attraction basin, the likelihood of the species falling into the survival attraction basin decreases, thereby increasing the risk of extinction.

### 9 Overview of Shared Socioeconomic Pathways

In response to the increasingly severe threat of global climate change to ecosystems, governments around the world have formulated and implemented a series of mitigation measures, such as the Paris Agreement, which seeks to limit global warming to well below 2°C above pre-industrial levels, and ideally to 1.5°C [17]. To achieve these goals, the scientific community has developed a framework known as Shared Socioeconomic Pathways (SSPs), indicating that future surface temperature increases are influenced not only by natural systems but also by human socioeconomic activities and development choices. SSPs outline five baseline scenarios: SSP1 (Sustainability–Taking the Green Road), SSP2 (Middle of the Road), SSP3 (Regional Rivalry–A Rocky Road), SSP4 (Inequality–A Road Divided), and SSP5 (Fossil-fueled Development–Taking the Highway) [18]. The relevant details are summarized in Table 1.

[7] Mark Newman. *Networks*. Oxford university press, 2018.

[8] Sergi Valverde, Jordi Piñero, Bernat Corominas-Murtra, Jose Montoya, Lucas Joppa, and Ricard Solé. The architecture of mutualistic networks as an evolutionary spandrel. *Nature Ecology & Evolution*, 2(1):94–99, 2018.

[9] Yamir Moreno, Romualdo Pastor-Satorras, Alexei Vázquez, and Alessandro Vespignani. Critical load and congestion instabilities in scale-freenetworks. *Europhysics Letters*, 62(2):292, 2003.

[10] Mercedes Pascual and Jennifer A Dunne. *Ecological networks: linking structure to dynamics in food webs.* Oxford University Press, 2005.
